# Accurate localization of cortical and subcortical sources of M/EEG signals by a convolutional neural network with a realistic head conductivity model: Validation with M/EEG simulation, evoked potentials, and invasive recordings

**DOI:** 10.1101/2024.04.30.591970

**Authors:** Hikaru Yokoyama, Natsuko Kaneko, Noboru Usuda, Tatsuya Kato, Khoo Hui Ming, Ryohei Fukuma, Satoru Oshino, Naoki Tani, Haruhiko Kishima, Takufumi Yanagisawa, Kimitaka Nakazawa

**Affiliations:** Institute of Engineering, Tokyo University of Agriculture and Technology, Koganei, Tokyo, 184-8588, Japan; Department of Life Sciences, Graduate School of Arts and Sciences, The University of Tokyo, Tokyo, 153-8902, Japan; Neural Prosthetics Project, Tokyo Metropolitan Institute of Medical Science, Setagaya, Tokyo, 156-8506, Japan; Sony Computer Science Laboratories. Inc., Tokyo, 141-0022, Japan; Japan Society for the Promotion of Science, Tokyo, 102-0083, Japan; Institute for Advanced Co-Creation Studies, Osaka University, Suita, Osaka, 565-0871, Japan; Department of Neurosurgery, Graduate School of Medicine, Osaka University, Suita, Osaka, 565-0871, Japan; Institute for Advanced Co-Creation Studies, Osaka University Graduate School of Medicine, Suita, Osaka, 565-0871, Japan

## Abstract

While electroencephalography (EEG) and magnetoencephalography (MEG) are well-established non-invasive methods in neuroscience and clinical medicine, they suffer from low spatial resolution. Particularly challenging is the accurate localization of subcortical sources of M/EEG, which remains a subject of debate. To address this issue, we propose a four-layered convolutional neural network (4LCNN) designed to precisely locate both cortical and subcortical source activity underlying M/EEG signals. The 4LCNN was trained using a vast dataset generated by forward M/EEG simulations based on a realistic head volume conductor model. The 4LCNN implicitly learns the characteristics of M/EEG and their sources from the training data without need for explicitly formulating and fine-tuning optimal priors, a common challenge in conventional M/EEG source imaging techniques. We evaluated the efficacy of the 4LCNN model on a validation dataset comprising forward M/EEG simulations and two types of real experimental data from humans: 1) somatosensory evoked potentials recorded by EEG, and 2) simultaneous recordings from invasive electrodes implanted in the brain and MEG signals. Our results demonstrate that the 4LCNN provides robust and superior estimation accuracy compared to conventional M/EEG source imaging methods, aligning well with established neuroscience knowledge. Notably, the accuracy of the subcortical regions was as accurate as that of the cortical regions. The 4LCNN method, as a data-driven approach, enables accurate source localization of M/EEG signals, including in subcortical regions, suggesting future contributions to various research endeavors such as contributions to the clinical diagnosis, understanding of the pathophysiology of various neuronal diseases and basic brain functions.

## 1. Introduction

Electroencephalography (EEG) and magnetoencephalography (MEG) are non-invasive brain imaging techniques that measure the electrical and magnetic fields generated by neuronal electrical activity. In contrast, other non-invasive measurements, like near-infrared spectroscopy (NIRS) and functional magnetic resonance imaging (fMRI), provide a low temporal resolution on the order of seconds due to their reliance on hemodynamic responses (Lindquist et al., 2009). M/EEG offers high temporal resolution on the millisecond order, directly reflecting the neuronal activity (Michel et al., 2004). Nevertheless, M/EEG’s spatial resolution is limited by the volume conduction effect (He et al., 2018). Enhancement of the spatial resolution of M/EEG will contribute to neurological applications, from diagnostics to understanding various brain functions.

Many initiatives have been undertaken to enhance the spatial resolution of M/EEG using electrophysiological source imaging techniques (ESI)(Sohrabpour and He, 2021). ESI represents an inverse problem that is inherently ill-posed, as there is no unique solution: multiple neural source combinations can produce identical M/EEG signal distributions (Nunez and Srinivasan, 2006). Traditional approaches require a priori assumptions, such as energy minimization or spatial smoothness, to deduce a unique solution (Hämäläinen and Ilmoniemi, 1994; Pascual-Marqui et al., 1994). However, the complexity of brain activity makes setting optimal regularization priors a significant challenge.

Recent advances in machine learning-based ESI techniques have sparked interest due to their data-driven approach that circumvents the need for explicitly defining a regularization term (Hecker et al., 2021; Jiao et al., 2022; Sun et al., 2022). These methods have demonstrated superior localization accuracy of estimated sources in cortical regions, compared to traditional ESI methods. The effectiveness of machine learning-based ESI in localizing subcortical sources remains to be confirmed.

Subcortical structures are pivotal to numerous brain functions, as highlighted in recent literature(Roy et al., 2022; Quian Quiroga, 2023). Their dysfunction is implicated in various diseases, such as Parkinson’s disease (Mirelman et al., 2019). Characterizing the electrical activity in subcortical regions is essential yet challenging due to the limitations of non-invasive techniques and the restrictions on invasive recordings to clinically relevant areas. Contrary to the long-held belief that deep brain structures are not discernible via M/EEG, recent studies have shown the potential of these techniques in detecting subcortical activity through simultaneous M/EEG and invasive electrode recordings (Pizzo et al., 2019; Seeber et al., 2019). These findings, along with the remarkable success of machine learning-based ESI methods in imaging cortical sources, suggest that such methods might also be adept at accurately imaging subcortical activities.

In this study, we explored the accuracy of ESI using a deep learning approach for both cortical and subcortical sources (Fig. 1). Based on the usefulness of convolutional neural networks (CNNs) in various types of inverse problems (Jin et al., 2017; Hecker et al., 2021; Crocker et al., 2023), we developed a 4-layer convolutional neural network (4LCNN) designed to estimate source activity from M/EEG topography, which was trained using a vast amount of data from forward M/EEG simulations based on a realistic head volume conductor model. The efficacy of our 4LCNN model for ESI was tested using a validation dataset created by forward M/EEG simulations and two types of human experimental data: (1) somatosensory evoked potentials (SEP) recorded by EEG, and (2) concurrent recordings from invasive brain electrodes and MEG signals.

**Figure 1.**
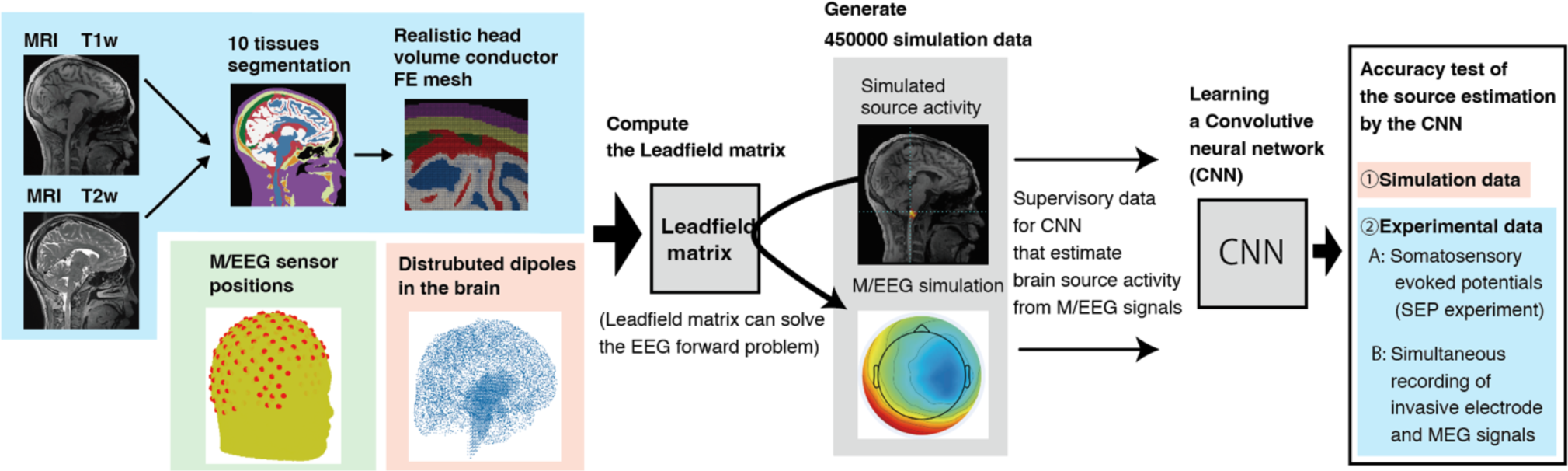
Conceptual diagram of this study. This flowchart shows how a M/EEG forward model is created (A), how a 4LCNN and forward M/EEG simulation were used to predict brain sources (B), and how the created 4LCNN model is validated (C).

## 2. Materials and Methods

### 2-1 Participants

For EEG data analysis, we obtained MRI and EEG data from three healthy adult males aged 23−25 years (Participant IDs: HY1−HY3). For MEG data analysis, MRI, MEG, Electrocorticographic (ECoG), and Stereo-electroencephalographic (SEEG) data were collected from three patients suffering from intractable epilepsy undergoing pre-surgical clinical assessment (Participant IDs: EP1−EP3, aged <u>10</u>’<u>s</u>−<u>20’s</u>). All participants provided written informed consent for participation in the study, which was conducted in accordance with the Declaration of Helsinki and approved by the Ethics Review Committee of The University of Tokyo (EEG experiment, reference number 701-2) and Osaka University (MEG experiment, reference number 14353). Structural MRI and M/EEG sensor position data were used for M/EEG simulation. M/EEG signals were utilized to evaluate source estimation for implementations with real data. ECoG and SEEG data were employed for the localization accuracy evaluation of estimated sources from the MEG signals.

### 2-2 EEG source estimation dataset

#### 2-2-1 EEG electrode location acquisition

The locations of EEG electrodes were digitized using an optical digitizing system (Brainsight, Rogue Research, Montreal, Canada). To achieve precise co-registration of MRI and EEG electrodes, we attached three donut-shaped multimodal (MR/CT) fiducial markers (PINPOINT, Berkley Corp., Bristol, CT) with a hole size of 1.27 mm onto the left and right pre-auricular points (LPA and RPA) and the forehead and digitized their locations. The coordinate systems of the electrode and MRI data were aligned based on these marker positions using the Fieldtrip software (Oostenveld et al., 2011).

#### 2-2-2 MRI Acquisition

T1 and T2-weighted structural MRIs were obtained using a Siemens Prisma 3T scanner (Siemens Medical Systems, Erlangen, Germany). As mentioned previously, three donut-shaped multimodal fiducial markers were attached to the head during MRI data acquisition for precise co-registration with EEG electrodes.

#### 2-2-3 Forward Model

We created forward models based on individual MRI data and digitized electrode locations to generate M/EEG data by forward simulation for generating training and validation datasets for 4LCNN. The relationship between the recorded potential by M/EEG sensors and brain source activity is expressed as follows:

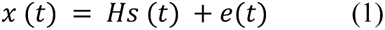

where *t* represents time, *x* (*t*) ∈ *R^n^*^×1^ is a vector of M/EEG signals measured by *n* sensors, H ∈ *R^n^*^×m^ is a leadfield matrix describing the electrical current flow from each dipole in the brain to every sensor, *s* (*t*) ∈ *R^m^*^×1^ represents the source signal generated by *m* brain sources, and *e*(*t*) ∈ *R^n^*^×1^ is the noise vector.

To simulate M/EEG data realistically and solve the inverse solution, we created a forward model based on anatomical head MRI images and M/EEG electrode positions.

We utilized SimNIBS equipped with the complete head anatomy reconstruction method (CHARM), which is a state-of-the-art technique, to automatically generate a highly accurate segmented MRI image with 10 tissues: white matter, gray matter, cerebrospinal fluid (CSF), scalp, eyeballs, compact bone, spongy bone, blood, muscle, and air pockets, based on individual T1 and T2 MRI images (Puonti et al., 2020). A hexahedral mesh was generated directly from the segmented MRI image using the ’prepare_mesh_hexahedral’ function within the Fieldtrip toolbox. To avoid staircase artifacts, we created geometry-adapted hexahedral meshes, where mesh nodes at tissue boundaries were slightly shifted to provide a smoother representation of the boundaries (Wolters et al., 2007). Then, utilizing finite element method (FEM) approaches, we constructed highly realistic multicompartment volume conductor models for forward EEG simulations. The conductivity (unit: S/m) of each tissue type was set as follows, based on a review article (McCann et al., 2019); white matter: 0.22, gray matter: 0.47, CSF: 1.71, scalp: 0.41, eyeballs: 0.5, compact bone: 0.0046, spongy bone: 0.050, blood: 0.57, muscle: 0.32. The conductivity for eyeballs was adopted from Haueisen et al., (1997) due to the absence of this information in McCann et al. (2019).

We employed T1 MRI data to obtain cortical surfaces and subcortical anatomical regions for defining the locations and orientations of the elementary dipole sources. The sources were placed on cortical surfaces and within subcortical volumes in accordance with a previous study (Krishnaswamy et al., 2017). Initially, cortical surfaces were extracted using FreeSurfer (Dale et al., 1999). Based on the cortical surface data, we created a high-resolution cortical surface triangular mesh with the HCP Workbench (Van Essen et al., 2013) and resampled it to approximately 8,000 sources using the iso2mesh toolbox (Tran et al., 2020). The edge lengths of the mesh were about 3 to 4 mm. Subcortical parcellations were obtained by FreeSurfer. The sources within subcortical volumes were evenly distributed with a grid resolution of 3 mm using the ’ft_prepare_sourcemodel’ function in Fieldtrip. For cortical sources, dipoles were placed with orientations normal to the cortical surface mesh (Samuelsson et al., 2021). For subcortical sources, triplets of orthogonal dipoles were situated within the subcortical volumes (Seeber et al., 2019). The leadfield matrix, which comprises vectors of potential amplitudes received by electrodes from each dipole source, was generated using the DUNEuro toolbox in Brainstorm (Schrader et al., 2021; Medani et al., 2023).

Furthermore, we developed a second forward model for validation of the 4LCNN. We modified the forward model used for training M/EEG data to create the second forward model for the validation, considering potential differences in brain-electric signal volume conduction between the actual head and our model, which cannot be precisely known a priori. We modified the electrode positions by adding Gaussian noise (mean: 0 mm, variance: 2 mm) to the three-dimensional coordinates of each electrode. Additionally, we altered the tissue conductivity in the FEM solution by scaling the original values by 1.2 or 0.8: white matter: 0.17 (original: 0.22), gray matter: 0.56 (original: 0.47), CSF: 1.36 (original: 1.71), scalp: 0.24 (original: 0.41), eyeballs: 0.6 (original: 0.5), compact bone: 0.0037 (original: 0.0046), spongy bone: 0.060 (original: 0.050), blood: 0.46 (original: 0.57), muscle: 0.38 (original: 0.32). All simulations in the validation section were performed using the validation model, whereas the calculation of the inverse solutions was based on the original model. The 4LCNN training data was simulated using only the original model. In this study, the original and modified models are referred to as the training model and validation models, respectively.

#### 2-2-4 EEG Simulations

We generated two different sets of simulated EEG signals: (1) EEG data for training the 4LCNN model using the training model and (2) validation data to test the constructed 4LCNN model against the validation model.

Each simulation involved a cluster of dipoles within each brain region for training the 4LCNN model and evaluating ESI methods (Samuelsson et al., 2021; An et al., 2022). For the simulation, cortical surfaces were divided into 62 cortical regions (31 regions per hemisphere) based on the DKT parcellation atlas (Klein and Tourville, 2012). Subcortical volumes were segmented into 17 anatomical regions (brain stem, left and right thalamus, caudate, putamen, pallidum, hippocampus, amygdala, accumbens area, and ventral diencephalon) as delineated by FreeSurfer segmentation. The cortical and subcortical segmentations were conducted using the recon-all pipeline in FreeSurfer. A source cluster for each region was constructed using the cortical mesh sheet and subcortical volume mesh described in the Forward Model section. Subcortical volume mesh was created from sources within the subcortical volumes using ‘delaunay’ and ‘triangulation’ functions in MATLAB. A single seed source was randomly selected; then, K nearest neighbors within the same brain region as the seed source were included in a cluster as active sources, where K was randomly chosen from 1 to 3. The number of dipoles in a cluster ranged approximately from 10 to 100. Dipole moments (amplitudes) were set to 1.0, 0.95, 0.9, 0.85 (in arbitrary units) for the seed and one, two, and three nearest neighbor dipoles, respectively. The orientations for cortical sources were constrained to be normal to the surface mesh, while subcortical source orientations were randomly determined, with a single orientation used within a cluster (Piastra et al., 2021). For the simulated data, Gaussian noise was added to the generated EEG signals for 159 electrodes. The noise level was adjusted to achieve a desired signal-to-noise ratio (SNR) in decibels (dB), defined as follows:

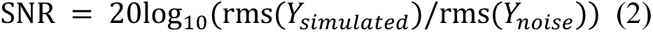

where *Y_simulated_* is a vector representing simulated EEG amplitude of the 159 channels EEG electrodes and *Y_noise_* is a vector representing added noise for the electrodes. The *rms(x)* is a function to calculate the root mean square of a vector *x*.

For training the 4LCNN model, we simulated 100,000 and 350,000 samples for cortical and subcortical sources, respectively, using the training model. For model evaluation, 100 samples were simulated for each cortical and subcortical source estimation using the validation model.

#### 2-2-5 Source Estimation by Convolution Neural Network

In this study, we utilized the 4LCNN to estimate source activity from M/EEG signals by addressing the M/EEG inverse problem. The 4LCNN model was specifically designed to estimate a single clustered source from the spatial distribution map of M/EEG signals.

As a preprocessing step for EEG signals, we standardized the amplitude of each spatial distribution map, comprising M/EEG signals from electrodes at a given time instance, to have a zero mean and unit variance. These maps were then transformed into input data for a 2D CNN by interpolating onto a 2D image of size 26 × 26. From this 2D matrix, the 4LCNN model estimates the activity of distributed dipoles within the brain. The design and training of the 4LCNN model were performed using MATLAB’s Deep Learning Toolbox on an NVIDIA RTX A6000 GPU.

The architecture of our 4LCNN model (Fig. 2) is based on ConvDip, a CNN model for cortical source estimation (Hecker et al., 2021). Our study aimed to estimate both cortical and subcortical source activity by enhancing the original CNN model to increase its expressive ability. We augmented the resolution of the input M/EEG map and the number of kernels (filters) in the CNN and expanded the convolutional layers beyond those in ConvDip. The first layer in the 4LCNN was a convolutional layer with 32 filters, each of size 3 × 3, employing zero-padding. Each convolutional layer was followed by a batch normalization (BN) layer (Ioffe and Szegedy, 2015) to accelerate training and mitigate internal covariance shift problems. A Rectified Linear Unit (ReLU) function followed each BN layer as the activation function. After four sets of convolution, BN, and ReLU layers, a dropout layer with a dropout rate of 30% was introduced to enhance the model’s generalizability. The final two layers of the 4LCNN were a fully connected layer and an output layer.

**Figure 2.**
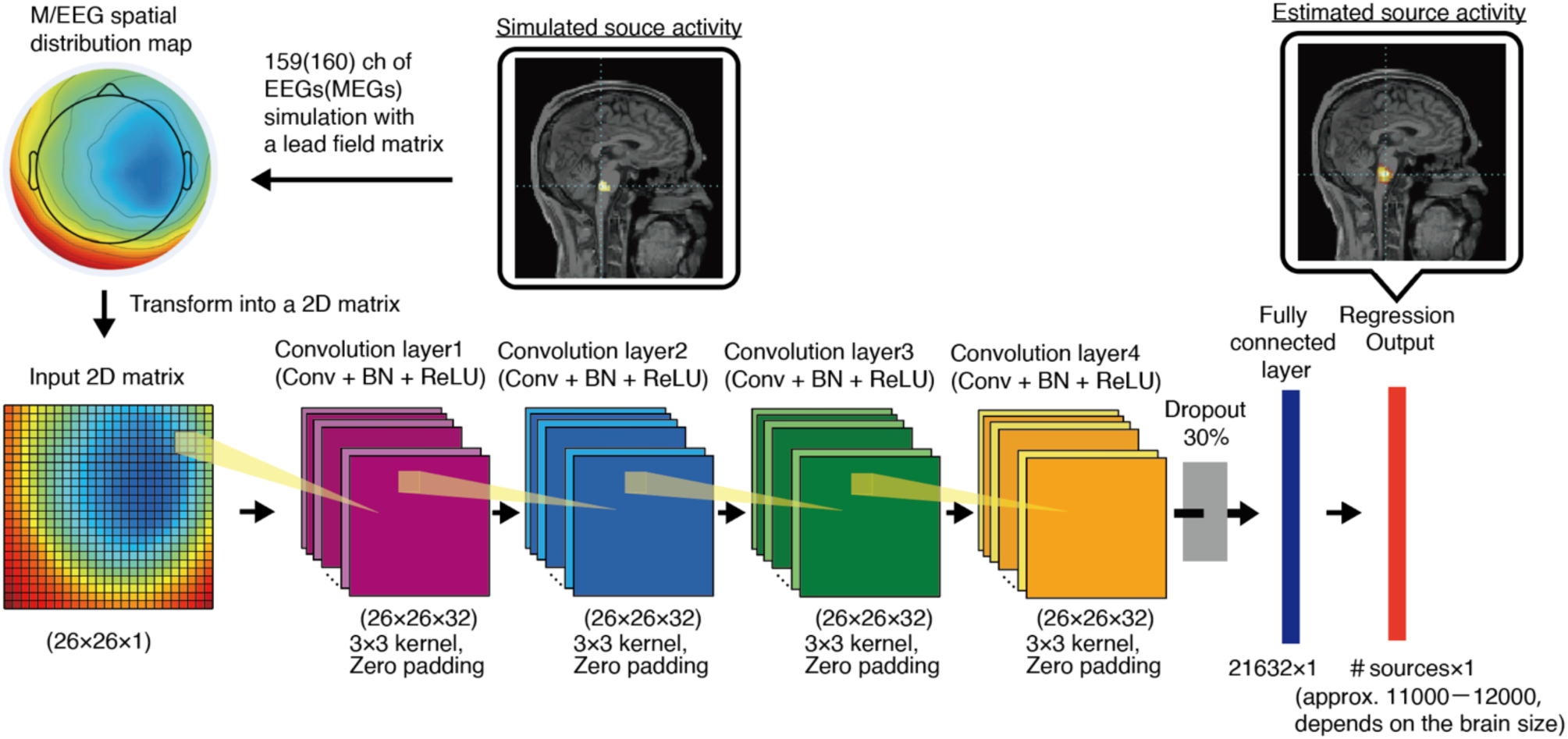
Architecture of the four layer convolutional neural network (4LCNN) to estimate source activity of M/EEG signals in the brain. The values from a simulated EEG data were interpolated to obtain a 26 × 26 matrix as an input (illustrated in the bottom left). The 2D matrix is sent to the subsequent four sets of convolution layers, which are consisted of convolution, batch normalization (BN) and Rectified Linear Unit (ReLU) sub-layers, have 32 convolution kernels of size 3 × 3 pixels with zero-padding. The convolution layers are followed by a dropout later (dropout rate: 30%) and fully connected (FC) layer consisting of 21632 neurons. Finally, the output layer contains neurons corresponding to activity of each source in the brain.

The 4LCNN model parameters were optimized using adaptive momentum estimation (ADAM) (Kingma and Ba, 2014) with default settings (learning rate = 0.001, β1 = 0.9, β2 = 0.999, ε = 1e-8). The loss function utilized was the root mean squared error (RMSE).

#### 2-2-6 Evaluation Metrics

We assessed the quality of inverse source estimation using three performance metrics:

I. **Localization error**: We calculated the Euclidean distance between the locations of the sources with the maximum value between the simulated and estimated data. This metric quantifies the accuracy of the estimated source center (Hecker et al., 2021).
II. **Spatial Dispersion:** To assess the spatial extent of sources, we calculated the Spatial Dispersion metric as follows (Hauk et al., 2022),

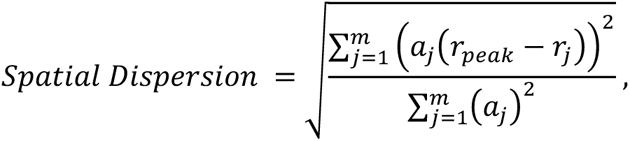

where, *m* is the number of sources, *a*_*j*_ and **r**_*j*_ are activity intensity and location of source *j*, respectively, 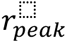 is location of source which has the maximum intensity across all sources, (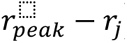) is the Euclid distance between 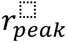 and *r_j_*.
III. **Area under the Precision-Recall curve (AUPRC)**: We computed the AUPRC to assess the accuracy considering both location and spatial extent of sources. The AUPRC is more suitable than other metrics, such as the Receiver Operating Characteristic (ROC) curve, for evaluating performance in imbalanced datasets like ours, which has a large imbalance between negatives and positives(Saito and Rehmsmeier, 2015). In our case, since simulated active sources are limited in a cluster, AU-PRC was considered to be a suitable method. The AUPRC quantifies the performance of a binary classification system as its discrimination threshold *T* is varied. The estimated sources are classified as active or inactive based on whether their amplitude exceeds the threshold *T*. We then tallied the true positives (TP), true negatives (TN), false positives (FP), and false negatives (FN) relative to the true sources. Precision and recall were computed as follows:

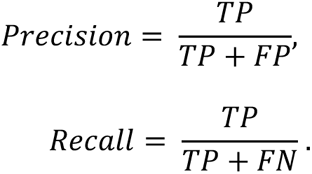

In the AUPRC analysis, precision and recall form the y-axis and x-axis, respectively. As *T* approaches 1, the number of positives approaches zero, leading to recall nearing zero, while precision becomes undefined (division by zero). At this limit, we used the precision value for the highest threshold below 1, assuming a horizontal asymptote as described in a prior study (Samuelsson et al., 2021). The PR curve was constructed by varying *T* from 0 to 1, and the AUPRC was integrated, yielding a value between 0 and 1. The closer the value to 1, the better the model’s performance.

#### 2-2-7 Experiments of Somatosensory Evoked Potentials

To validate the 4LCNN model with real EEG data, we employed SEP experiments due to the accumulation of evidence supporting the neural origin of SEP components. EEG and depth electrode recordings have confirmed that somatosensory information from median nerve stimulation propagates sequentially through the cervical spinal cord, medulla, thalamus, and somatosensory cortex at approximate intervals of 11 ms, 13‒14 ms, 16‒17 ms, and 20 ms, respectively (Macerollo et al., 2018). We administered electrical monophasic square wave constant current pulses for a duration of 200 ms from a clinical constant current stimulator (DS7A, Digitimer Ltd., Welwyn Garden City, United Kingdom) to stimulate the right median nerve. The electrode pair was placed on the wrist of the right hand. The current intensity was set to the sum of the motor and sensory thresholds for each participant, in line with the International Federation of Clinical Neurophysiology’s recommendations (Nuwer et al., 1994). Stimulation was applied 2000 times at a repetition rate of 4.98 Hz, with the entire session repeated four times, totaling 8000 stimulations. To maintain the participants’ attention to the stimulus, pauses were included approximately every 45– 90 seconds. During each session, stimulation was paused randomly for about 2 seconds between 4 to 7 times. The duration of each session was approximately 7 minutes, pauses included. Participants were instructed to count the number of pauses and report their count at the session’s end. All participants accurately reported the number of pauses for all sessions.

EEG signals were recorded from 159 channels using an EEG amplifier (ActiCHamp Plus, Brain Products, Germany) with active electrodes (ActiCap Slim, Brain Products GmbH, Germany) at a sampling rate of 5000 Hz during the SEP experiment. Electrode impedances were maintained below 10 kΩ, significantly lower than the recommended impedance (<50 kΩ) for high-impedance EEG amplifiers. The same three-dimensional electrode coordinates employed for the head model creation were used, as the electrode position data was measured immediately prior to the SEP experiment. The EEG signals were band-pass filtered with a 4th order Butterworth filter at a cutoff frequency of 100–250 Hz (Buchner et al., 1995; Fukuda et al., 2008). This high-frequency component was selected to minimize baseline fluctuation and to separately evaluate components occurring closely at 13–14, 16–17, and 20 ms, corresponding to the medulla, thalamus, and primary somatosensory cortex components, respectively. SEPs from all participants exhibited distinct peaks corresponding to these components. We estimated the source activity of these components from the EEG spatial distribution map at the peak times using the 4LCNN model.

### 2-3 Validation with simultaneous recording of MEGs, SEEG and ECoG data

To further evaluate the 4LCNN model on the real data, we conducted ESI using the 4LCNN on MEG data and evaluated its accuracy with simultaneously recorded invasive brain signal recordings from SEEG and ECoG (Fig. 3). This study included three patients diagnosed with epilepsy.

**Figure 3.**
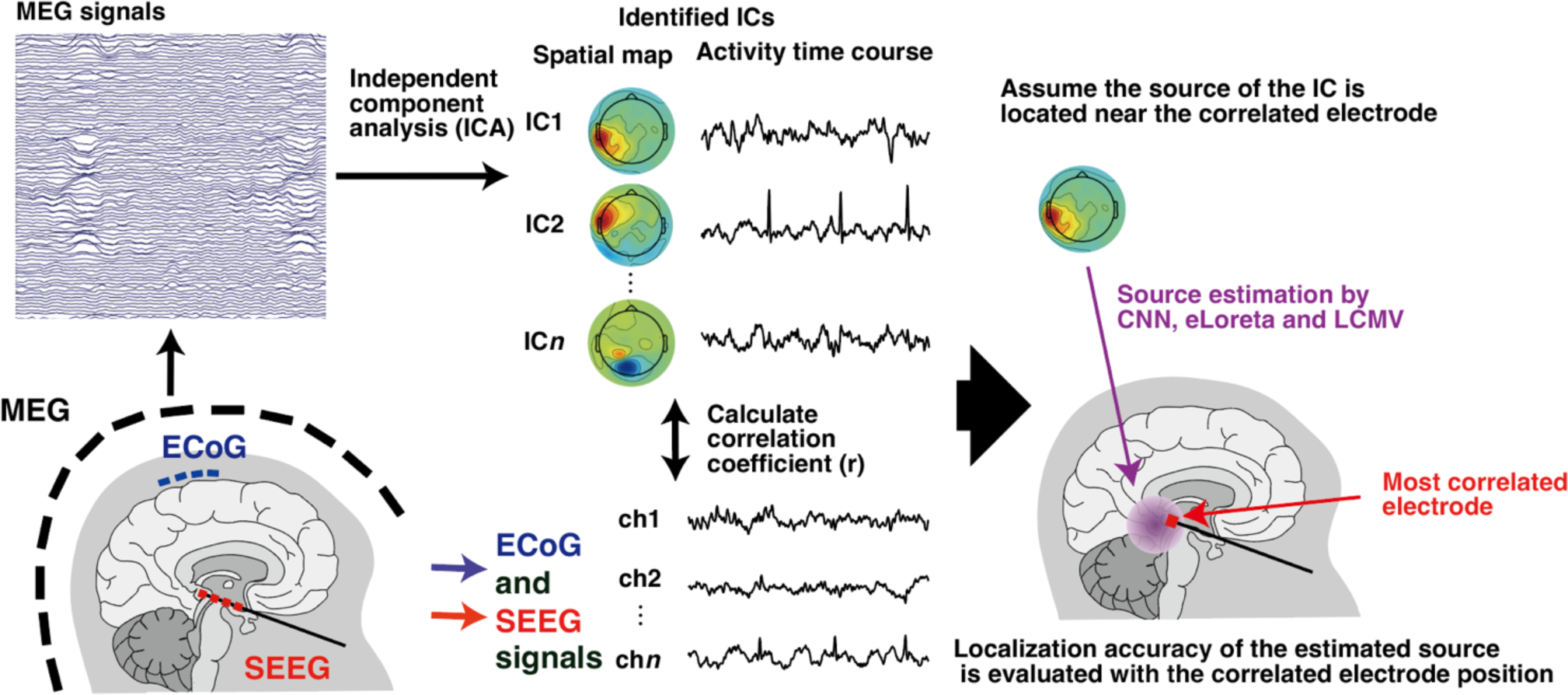
Conceptual diagram of the ESI method validation with simultaneous recordings of magnetoencephalographic (MEG), electrocorticographic (ECoG) and stereo-electroencephalographic (SEEG) signals. The MEG signals were decomposed into independent components by independent component analysis (ICA). Similarity of time courses of the ICs and intracranial signals were evaluated by the correlation coefficients. Then, Assuming the source of the IC is located near the correlated electrode, we tested localization accuracy of estimated sources and corresponding intracranial electrodes.

#### 2-3-1 Data acquisition

We simultaneously recorded MEG, SEEG, and ECoG signals from the participants in a resting state (with eyes closed, relaxed, possibly asleep) for 5 minutes. This recording was repeated four to six times per participant, accumulating 20-30 minutes of data for each participant.

MEG data were recorded using a 160-channel whole-head MEG system (MEG Vision NEO, Yokogawa Electric Corporation, Tokyo, Japan). Participants lay supine in a magnetically shielded room with five head-marker coils attached to their face. The positions of these head-marker coils were measured before and after each MEG recording to determine the relative position and orientation of the MEG sensors to the participant’s head. The MEG signals were sampled at 2000 Hz with a band-pass filter setting of 0.1−500 Hz. To co-register MEG data with individual MRI data, the three-dimensional locations of 50 points on each participant’s scalp and facial surface were digitized (FastSCAN, Polhemus, Colchester, Vermont, USA).

SEEG and ECoG electrodes were implanted and their locations in each participant are shown in Figs. S1−S3. We collected data from 24–54 SEEG contacts and 42–70 ECoG contacts, recorded at 2000 Hz by an digital EEG system (EEG-1200, Nihon Koden, Tokyo, Japan). All subcortical electrodes were referenced to average potential of two electrodes placed subcutaneously or on the cortex during the recordings. Initially, signals were filtered using a band-pass filter ranging from 1-60 Hz,.

Pre- and post-implantation T1-weighted MRI scans were conducted with Discovery MR750 (GE Healthcare, Chicago, U.S.A) for EP1 and SIGNA Architect (GE Healthcare) for EP2 and EP3.

MRI data were anonymized in accordance with the ethical policy for sharing data from the measuring organization, Osaka University, to other joint research institutes. The AnonyMI algorithm (Mikulan et al., 2021) was employed for anonymization to significantly reduce the risk of re-identification while conserving geometrical information. After electrode implantation, a CT scan was also performed to ascertain the positions of the implanted electrodes with the Aquilion Precision scanner (Toshiba Medical Systems, Tokyo, Japan).

#### 2-3-2 Data analysis

We created 4LCNN models for estimating MEG source activity using the same methodology as for EEG source estimation, with minor modifications tailored for MEG. The spatial alignment of MEG sensors and MRI data was based on the coordinates of the head surfaces digitized during the MEG experiment and the head surfaces extracted from T1-weighted MRI images. This alignment was achieved through rigid-body transformations to minimize the distance of these points to the head surface using the Brainstorm software. MRI and CT data were also aligned using the ’ft_convert_coordsys’ function in Fieldtrip. ECoG and SEEG electrode locations were then determined using MRI and CT data via a pipeline designed for merging anatomical images with intracranial electrode placement in Fieldtrip (Stolk et al., 2018).

MEG signals were band-pass filtered within a frequency range of 1 to 60Hz, and then re-referenced with a common average reference (CAR) technique (Fahimi Hnazaee et al., 2020). Subsequently, independent component analysis (ICA) was performed using the ’runica’ function in the EEGLAB toolbox (Delorme and Makeig, 2004). Correlation coefficients (*r*) between the time courses of the independent components (ICs) and the signals from SEEG and ECoG electrodes were calculated. For each IC, if *r* values exceeded 0.2 for any electrode, the electrode with the highest *r* was assumed to be most proximal to the source of the IC. We then computed the Euclidean distance between the estimated location of the IC and the corresponding electrode location.

### 2-4 Comparisons to other ESI methods

We compared the performance of the 4LCNN model with ConvDip (Hecker et al., 2021), a previously published CNN model for ESI, and two commonly utilized ESI methods—eLORETA (Pascual-Marqui et al., 2011) and the LCMV beamformer (Van Veen et al., 1997)—using the same set of simulations and experimental data. ConvDip was trained with the same forward M/EEG simulation data as the 4LCNN using MATLAB’s Deep Learning Toolbox. The eLORETA and LCMV analyses were conducted using Fieldtrip software. Both eLORETA and LCMV require noise covariance information among sensors. We utilized an identity matrix for the noise covariance for the simulated EEG data because white Gaussian noise was used in the EEG simulation. For MEG ICs, an identity matrix was also employed since the noise covariance among sensors within a single IC was indeterminate, following a recent study (Michalareas et al., 2023). For SEP data, covariance matrices were derived from signals during the pre-stimulation period (-50 ms to -5 ms) in accordance with previous studies on evoked responses (Dmochowski et al., 2015; Hecker et al., 2021). The regularization parameter for the noise covariance matrix in eLORETA and LCMV was set to SNR^-1^ for the simulation and SEP data (Samuelsson et al., 2021), and a default value of 0.05, as per Fieldtrip’s settings, was used for the single ICs of MEG.

### 2-5 Statistics

The normality of the datasets was evaluated using the Lilliefors test. For all ESI methods, estimated activity was normalized to the peak value across sources. With respect to the three performance metrics—localization error, spatial dispersion, and AUPRC—for the accuracy validation with simulation data, the differences between the ESI methods were statistically compared using a permutation test, a non-parametric method (Nichols and Holmes, 2001), across each SNR condition and metric. The permutation test was selected because the normality was not observed in all cases. P-values derived from the permutation test were adjusted for multiple comparisons using the false discovery rate (FDR) control method (Benjamini and Hochberg, 1995).

We also explored the relationships of activation areas between the simulated and estimated data in the simulation dataset. The activation area was defined as the number of active sources with a value exceeding 0.5.

In the ESI analysis of SEP data from healthy participants, the spatial dispersion metric was computed, as this metric can be determined without true source activity information. Spatial dispersion was compared among the ESI methods using a paired t-test with FDR correction since normality was confirmed.

In the ESI analysis of resting state MEG data from individuals with epilepsy, both localization error and spatial dispersion were compared among ESI methods using permutation tests with FDR correction due to the lack of normality in the datasets.

## 3. Results

### 3-1. Simulation data

Figure 4 shows examples of estimated cortical source activity by 4LCNN, ConvDip, LCMV, and eLORETA. The simulated sources were located in the medial part of the right superior frontal gyrus (Fig. 4A) and the left middle temporal cortex (Fig. 4B). Visual inspection suggests that both 4LCNN and ConvDip accurately estimated the location and spatial spread, even at a low SNR condition (SNR = 10 dB). Conversely, the LCMV generally accurately estimated the location of activity but exhibited large spatial spreads. Variations in noise level did not seem to cause significant changes. eLORETA’s estimated activities were dispersed at an SNR of 10 dB, but at higher SNR conditions, the estimations appeared more focal and accurate. However, eLORETA and LCMV failed to differentiate the left and right sides of the medial part of the superior frontal gyrus separately (Fig. 4A).

**Figure 4.**
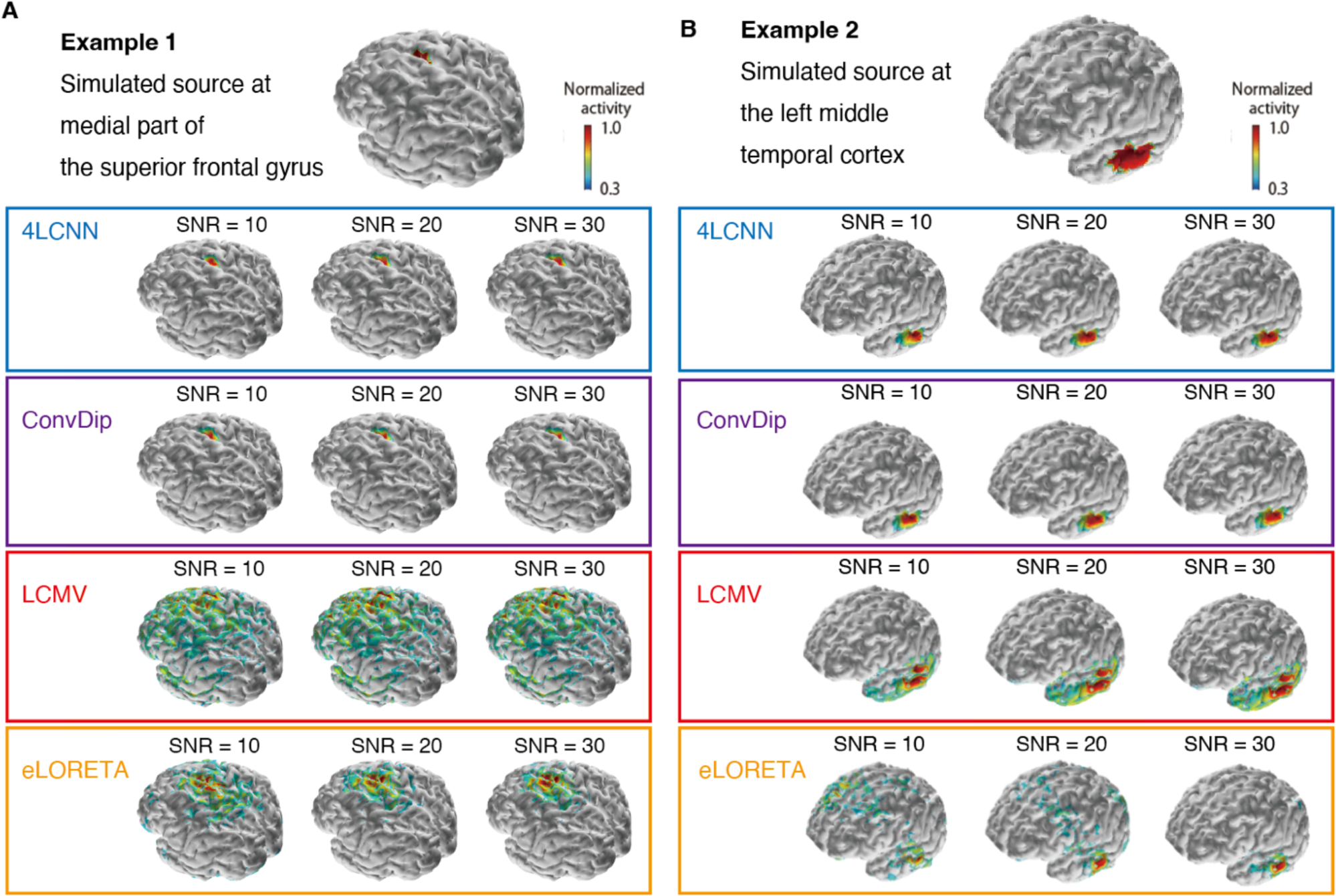
Examples of estimated source activity of cortical sources by 4LCNN, ConvDip, LCMV and eLORETA. Examples of estimated sources from EEG for the simulated sources in the medial part of the right superior frontal gyrus (A) and the left middle temporal cortex (B). Different noise levels were added to EEG simulations, and estimation results for three different SNR conditions (SNR = 10, 20, and 30 dB) are shown.

Figures 5 and S4 present examples of estimated subcortical source activity. The simulated sources were located in the right thalamus (Fig. 5A) and the left pallidum (Fig. S4A). The 4LCNN method accurately estimated the location and spatial spread of these examples (Figs. 5B and S4B), with no activity observed in the cortical regions. This accurate estimation was consistent across SNR conditions, with the exception of the left pallidum at a low SNR condition (SNR = 10 dB, Fig. S4B). ConvDip estimated activities near the subcortical structures without cortical activity for both the right thalamus and the left pallidum (Fig. 5C and S4C). However, the maximum activity was erroneously estimated in adjacent structures that were not the targets. LCMV estimated maximum activities in the cortical regions for both the right thalamus (Fig. 5D) and left pallidum data (Fig. S4D). For the right thalamus simulation (Fig. 5D), while broad activities were estimated within deep brain regions, the maximum activity was incorrectly localized in the medial orbitofrontal or insular cortices. In the pallidum simulation (Fig. S4D), the maximum activity was wrongly estimated in the left medial temporal lobe, with high activity scattered across several subcortical regions. Similar to the cortical source simulation examples (Fig. 4), the SNR condition did not significantly impact LCMV estimation. For eLORETA (Figs. 5E and S4E), in low SNR conditions, estimated activities were widely scattered across cortical and subcortical regions, with maximum values mistakenly placed in the cortical regions for the examples of the right thalamus and the left pallidum. At a high SNR condition (SNR = 30), eLORETA accurately located the maximum activity for both examples but the estimated activities were broadly distributed over several subcortical and medial cortical regions (Figs. 5E and S4E).

**Figure 5.**
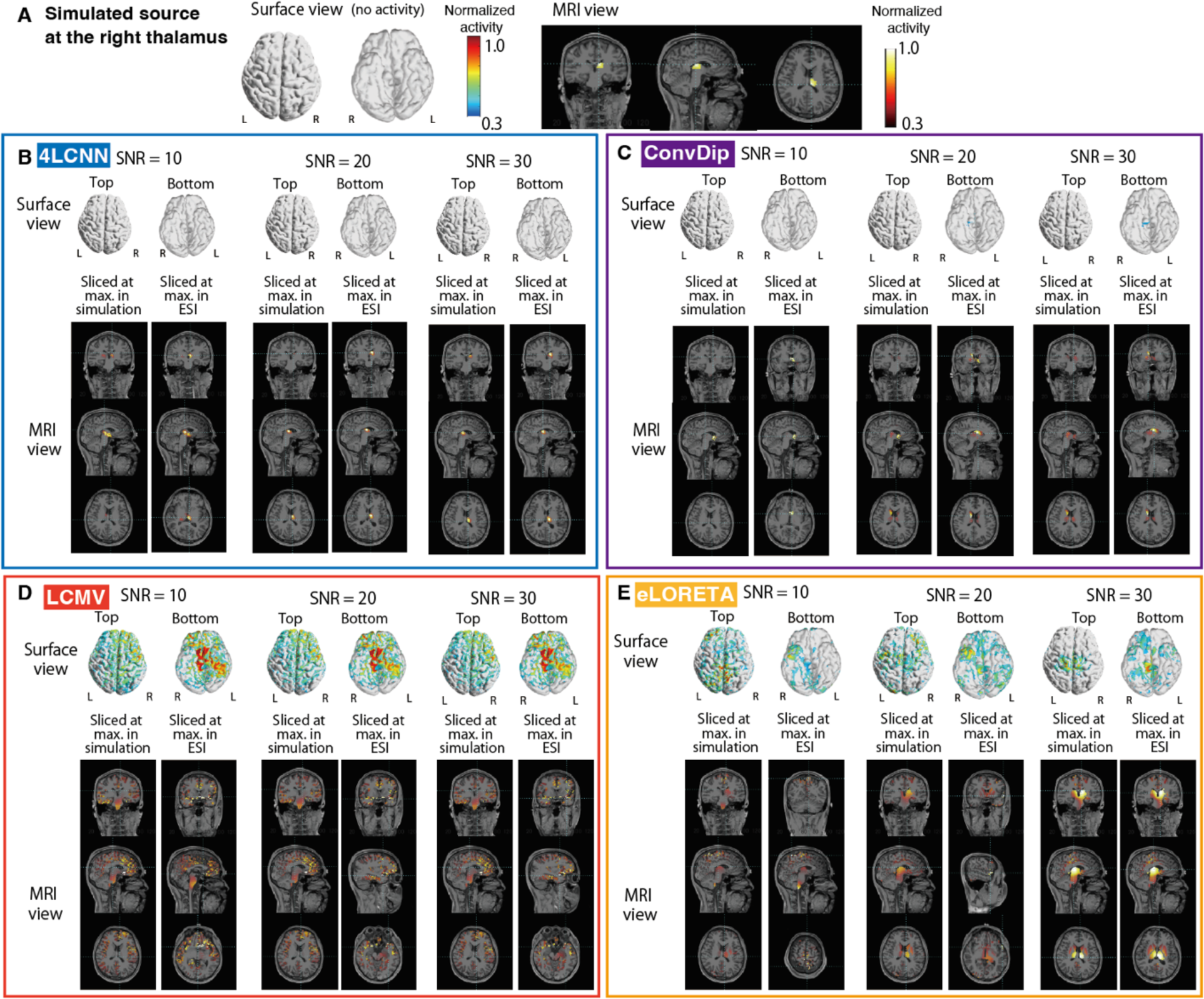
Examples of estimated subcortical source activity located at the right thalamus. (A): Simulated source activity is shown in surface and MRI views, with no activity in the surface view as this example pertains to a hippocampal source. (B-E): Estimated source activity by 4LCNN, ConvDip, LCMV, and eLORETA are presented in (B), (C), (D), and (E), respectively. EEG simulations incorporated various noise levels. Estimation results for three different SNR conditions (SNR = 10, 20, and 30 dB) are depicted. Activities are displayed in surface and MRI views. MRI views are sliced at two different locations: one at sources with the maximum values in the simulated sources, and another at the maximum values according to the targeted ESI method.

We calculated three metrics (localization error, spatial dispersion and AUPRC) to quantitatively evaluate the ESI performance of the 4LCNN compared to the other methods (Fig. 6). The localization error of the 4LCNN was significantly lower than that of the other ESI methods across all SNR conditions for both cortical and subcortical sources (Fig. 6A) (p < 0.05, see Supplemental Excel data file for detailed statistical values). For cortical sources, the mean error was below 10 mm in the 4LCNN at SNRs higher than 10 dB, reaching to 6.6 mm at an SNR of 30 dB. ConvDip showed relatively low localization errors; however, eLORETA was more accurate than ConvDip in high SNR conditions. Despite this, the mean error for eLORETA was 9.3 mm at an SNR of 30 dB.

**Figure 6.**
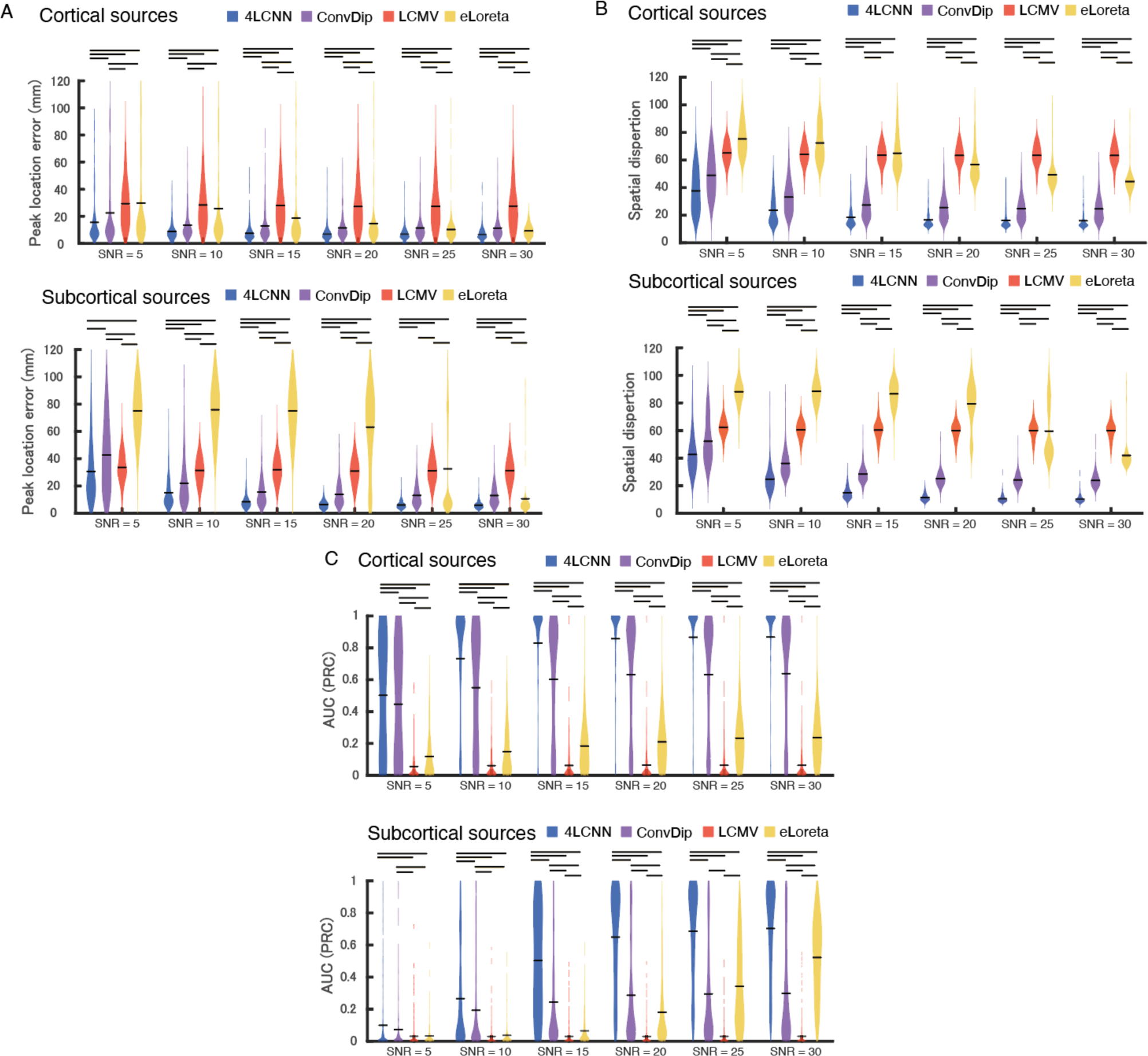
Quantitative evaluation of estimation accuracy in EEG simulation data. Violin plots illustrate the distribution of three different metrics (localization error of peak activity, spatial dispersion, and area under the Precision-Recall curve (AUPRC)) based on 300 simulation data points from three participant head models (one hundred for each) in (A), (B), and (C), respectively. The horizontal bars in the violin plots indicate the mean values. Bars above the violin plots denote significant differences between conditions (p < 0.05, according to the permutation test with FDR correction).

For subcortical sources, 4LCNN also showed superior performance regarding localization error compared to other methods (p < 0.05, see Supplemental Excel data file for detailed statistics) except at an SNR of 5 dB, where no significant difference from LCMV was found. The mean error in the 4LCNN was below 10 mm at SNRs higher than 15 dB and decreased to 5.9 mm at an SNR of 30 dB. ConvDip generally had lower localization errors than LCMV and eLORETA; however, eLORETA was more accurate than ConvDip at an SNR of 30 dB. On the other hand, at SNRs below 25 dB, the error for eLORETA was larger than LCMV and ConvDip.

Spatial dispersion was consistently lower in the 4LCNN compared to the other ESI methods in all conditions for both cortical and subcortical sources (Fig. 6B) (p < 0.05, see Supplemental Excel data file for detailed statistics). The 4LCNN tended to estimate source activity more focally than the other two ESI methods.

The AUPRC, a comprehensive metric assessing the central location and spatial extent of estimated sources, was significantly higher in the 4LCNN than the other ESI methods across all SNR conditions for both cortical and subcortical sources (Fig. 6C) (p < 0.05, see Supplemental Excel data file for detailed statistics).

These results (Fig. 6) were derived from 300 simulations based on head models from three participants for validation (one hundred for each). Figure S5 displays all three metrics for each participant’s head model, showing consistent results with the pooled data in Figure 6.

We then examined the relationship between the simulated and estimated activation area sizes (Fig. 7). As SNR increased, a stronger positive correlation between simulated and estimated area sizes was observed with the 4LCNN. For cortical sources (Fig. 7A), a moderate to high positive correlation was noted (r = 0.43 to 0.68) at SNRs of 15 or above. For subcortical sources (Fig. 7B), weak to moderate positive correlations were observed (r = 0.32 to 0.41) at SNRs of 15 or above. In contrast, negligible to very weak negative correlations were observed (r = -0.19 to 0.14) with the other ESI methods for both cortical and subcortical sources.

**Figure 7.**
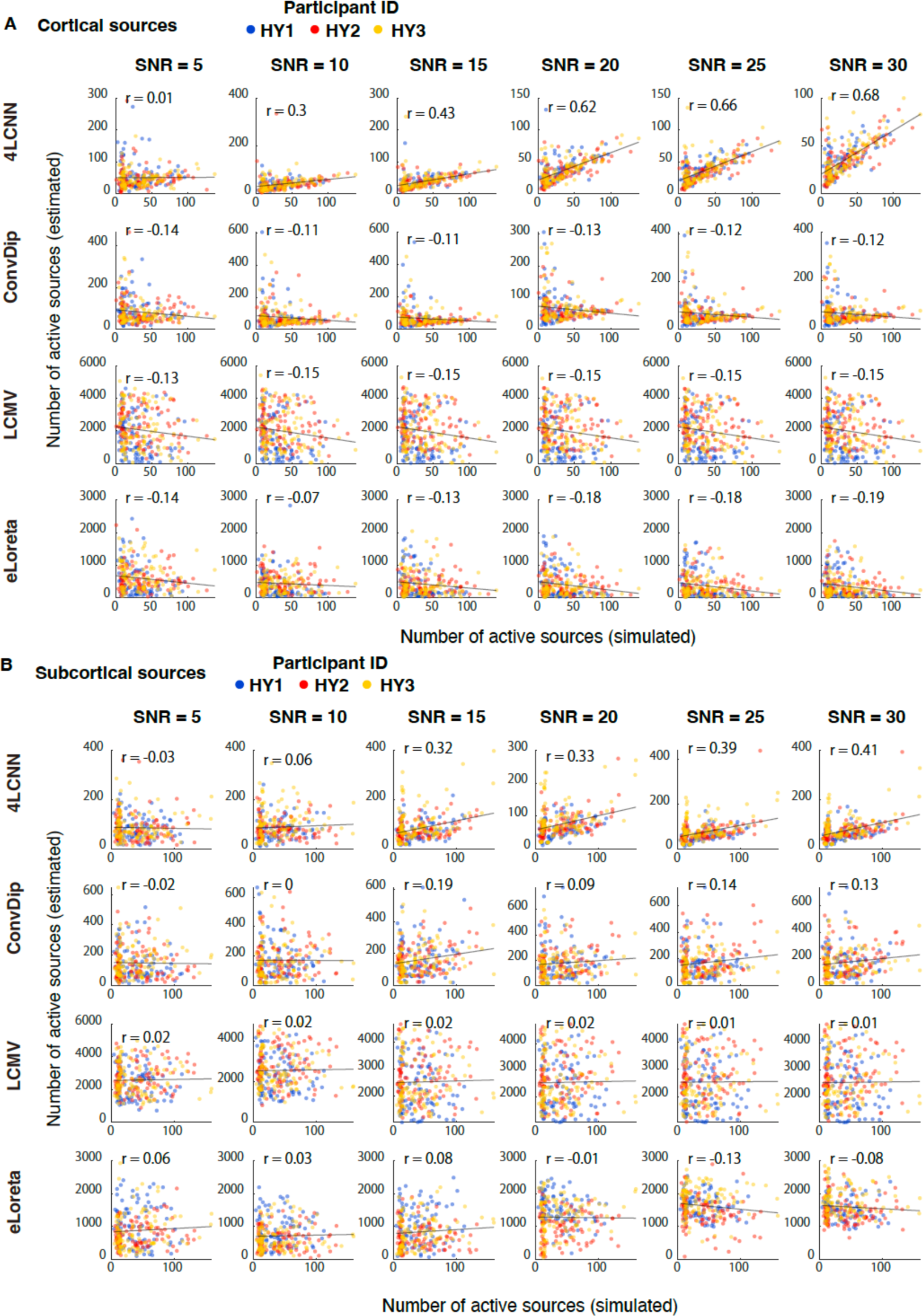
Correlation of activation area (i.e., number of active sources) between simulated and estimated source activities of 300 simulation data from three participants’ head models for cortical source simulation (A) and subcortical source simulation data(B). Each participant data is plotted as closed circles in a different color.

### 3-2. SEP data

In addition to the simulation data, we further validated the 4LCNN model using real human data. For this purpose, SEP data acquired from healthy participants was utilized. It is hypothesized that somatosensory information from median nerve stimulation travels through the cervical spinal cord, medulla, thalamus, and somatosensory cortex at approximately 11 ms, 13-14 ms, 16-17 ms, and 20 ms, respectively (Fig. 7A) (Macerollo et al., Trends Neurosci., 2018). SEP data from all participants exhibited clear peaks corresponding to the SEP components (Figs. 8A, 8C, and S6). From the SEP components, we focused on the latter three as they originate from the brain. Each SEP component presented distinct topographic maps (Fig. 7D), and these maps were consistent across participants (Figs. 7D and S6).

**Figure 8.**
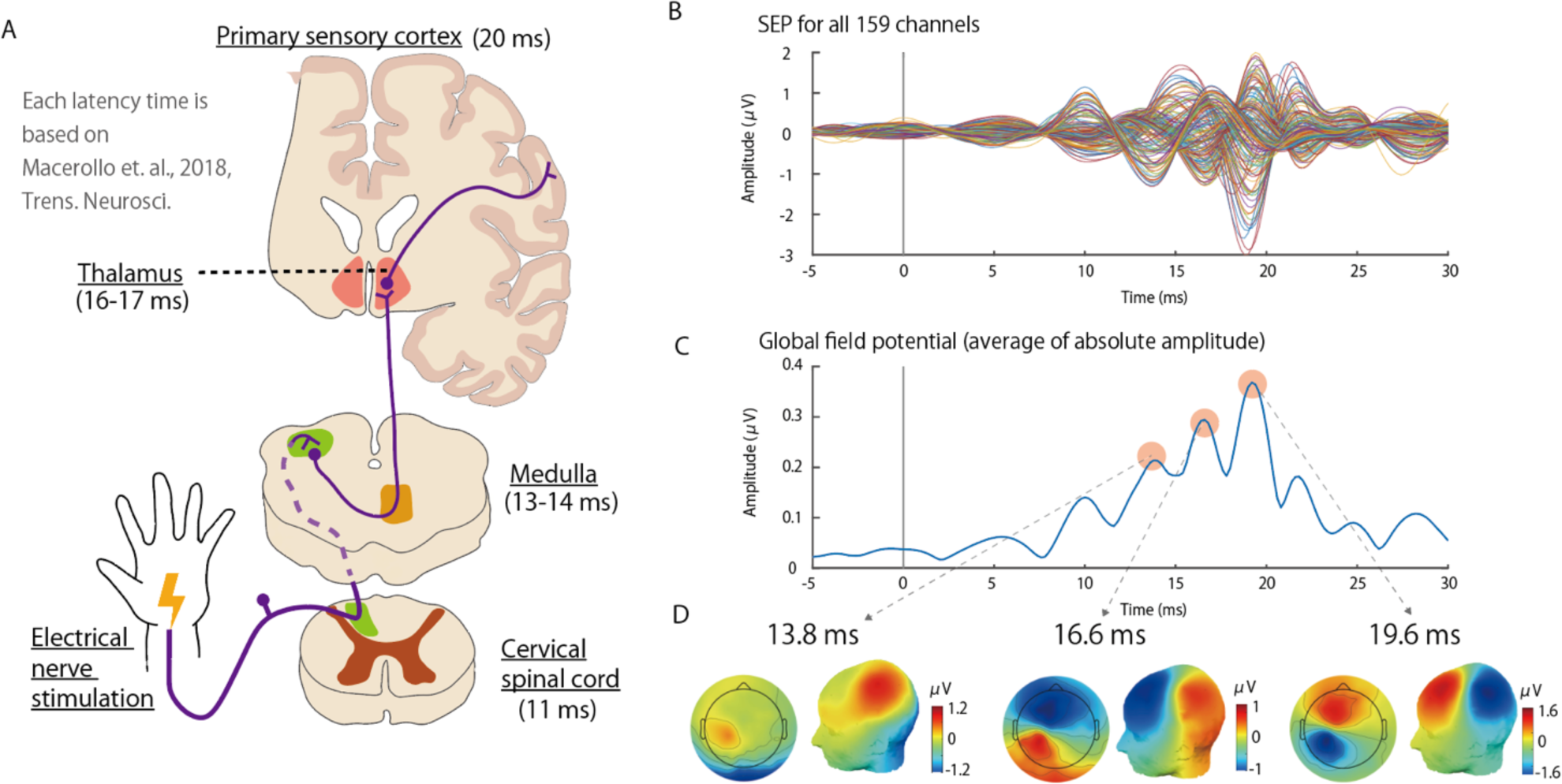
Somatosensory evoked potentials (SEPs) in a participant (participant ID: HY1). (A): A diagram outlining the presumed neural pathway and latencies for SEP. (B): An example of SEPs recorded across all 159 channels. (C): Global field potentials, representing the mean absolute amplitude across all channels, showing three distinct peaks corresponding to the components depicted in (A). (D): Topographic maps for the peak SEP activities, presented in both 2D and 3D.

We estimated source activity based on the topographic distribution of amplitudes from SEP components. The estimation results for participants HY1, HY2, and HY3 are shown in Figs. 9, S7, and S8, respectively. Using the 4LCNN method (Figs. 9B, S7B, and S8B), the 13 ms component’s activity was accurately localized in the medulla for all participants, aligning with assumptions. For the 17 ms component, activity was localized in the left thalamus for two participants (HY1 and HY2). For the third participant (HY3), although the maximum activity was found slightly above the left thalamus in the left caudate, significant activity was also present in the left thalamus. The 20 ms component was consistently localized in the hand region of the left primary somatosensory cortex for all participants. In summary, the estimated source activity of SEP components was generally in accordance with a neural pathway of the somatosensory system (Cruccu et al, 2008).

**Figure 9.**
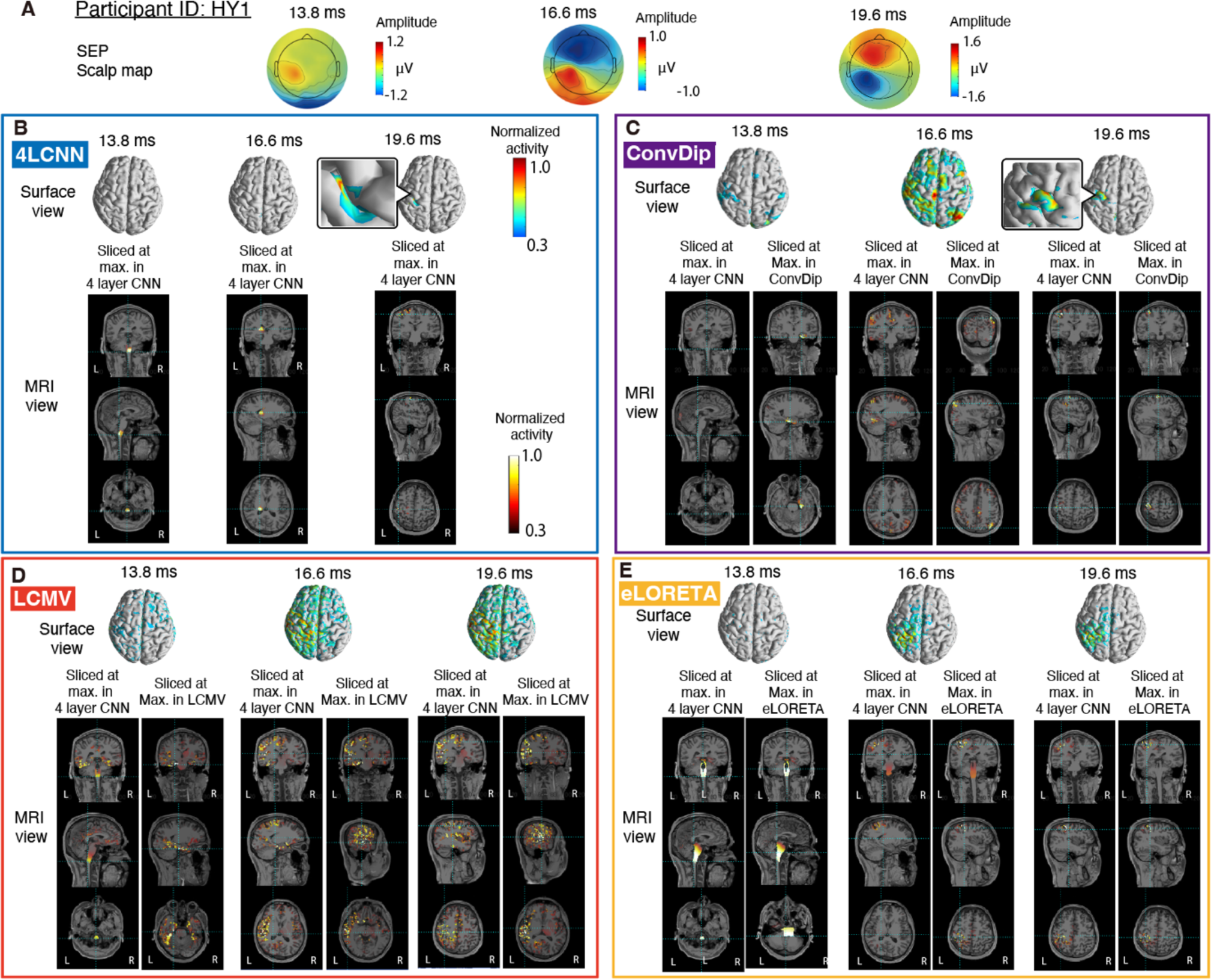
Source activity estimation of SEP in a participant (participant ID: HY1). (A): Topographic maps depicting three peak SEP activities. (B-E): Estimated source activities are presented in surface and MRI views, with (B-E) displaying estimations by 4LCNN, ConvDip, LCMV, and eLORETA, respectively. MRI views are sliced at sources with the maximum values in 4LCNN and the target ESI method respectively.

ConvDip (Figs. 9C, S7C, and S8C) estimated the maximum activity for the 13 ms component in the mesial lobe (HY1) and the brainstem (HY2 and HY3). For the 17 ms component, the maximum activity was misplaced in the right parietal cortex (HY1), the left amygdala (HY2) and left ventral diencephalon (HY3), rather than the left thalamus. Estimated activity for the 20 ms component was observed in the left primary sensory area (HY1–HY3) and additionally in the left primary motor area for two participants (HY1 and HY3).

LCMV (Figs. 9D, S7D, and S8D) incorrectly localized the maximum activity for the 13 ms component in the mesial lobe for all three participants, with either large (HY1) or minimal (HY2 and HY3) medullary activity. All participants showed activity across extensive areas of the cerebral cortex for the 17 ms component, with pronounced activity in the left sensorimotor area, mesial lobe, and temporal lobe. The estimated activity for the 20 ms component was similar to that for the 17 ms component.

eLORETA (Figs. 9E, S7E, and S8E) showed weak (HY2) or strong (HY1 and HY3) activity in the brainstem for the 13 ms component, yet only HY1 correctly localized the maximum activity in the brainstem (pons, not medulla). The other two (HY2 and HY3) exhibited maximum activity in the temporal (HY2) and occipital cortices (HY3). For the 17 ms component, despite observing weak activity in the left thalamus (HY3, HY2), eLORETA incorrectly estimated large activity areas in the left sensorimotor area. For the 20 ms component, the primary activity was noted in the left primary sensory area; however, unlike with 4LCNN, there was no distinct peak activity localized in the hand sensory region.

We also analyzed the spatial dispersion of the estimated source activity from SEP components (Fig. 10). As with the simulation data (Fig. 6C), spatial dispersion for the 4LCNN was significantly lower than that observed in the other ESI methods (4LCNN vs. ConvDip: p = 0.0083, effect size (ES) = 0.43; 4LCNN vs. LCMV: p = 0.0032, ES = 1.97; 4LCNN vs. eLORETA: p < 0.001, ES = 1.61; ConvDip vs. LCMV: p < 0.001, ES = 2.01; ConvDip vs. eLORETA: p = 0.020, ES = 1.46; LCMV vs. eLORETA: p = 0.76, ES = 0.14).

**Figure 10.**
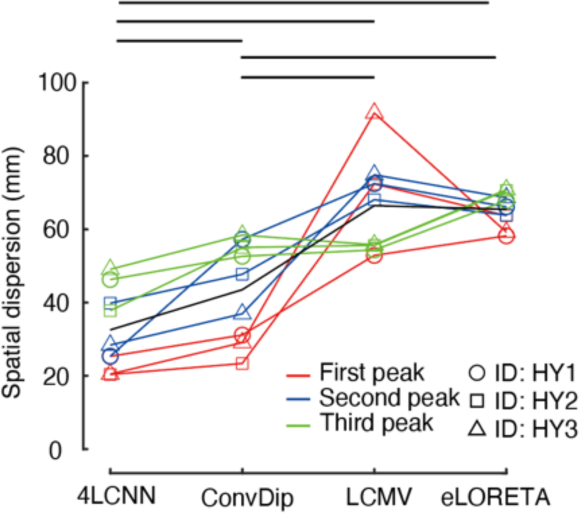
Spatial dispersion of estimated source activities from SEP signals. Comparisons of spatial dispersion values among different ESI methods are shown. The horizontal bars indicate significant differences (p < 0.05, paired t-test with FDR correction). Data corresponding to each participant and SEP peak timing are differentiated with various markers and colors.

### 3-3. MEG, ECoG, SEEG data

In addition to SEP data, the validity of the 4LCNN model was tested using a dataset from simultaneous recordings of MEG, ECoG, and SEEG signals from three individuals with epilepsy. Prior to this validation, the accuracy of the ESI methods was verified using MEG simulation data (Fig. S9), which yielded results similar to those obtained with EEG simulation data (Fig. 6).

Source activities of ICs separated from MEG signals were estimated. If the activity time series of the ICs correlated with signals from invasive electrodes, the Euclidean distance between the location with the maximum activity in the estimated source activity and the most correlated electrode was calculated. Examples from a participant (ID: EP1) are provided in Fig. 11. Activation time series of two ICs were correlated electrodes located at the temporal lobe (i.e., an ECoG electrode) and the hippocampus (i.e., a SEEG electrode), respectively. the topography maps of two correlated ICs are shown in Figs. 11A and 11B. Distances between the locations of maximum activity and the most correlated electrodes in the temporal lobe and hippocampus ranged from 7.3 to 38.5 mm for all ESI methods (Fig. 11). Notably, in this example, the 4LCNN method estimated a source location within the hippocampus that was very close to the corresponding electrode (7.3 mm, Fig. 11B).

**Figure 11.**
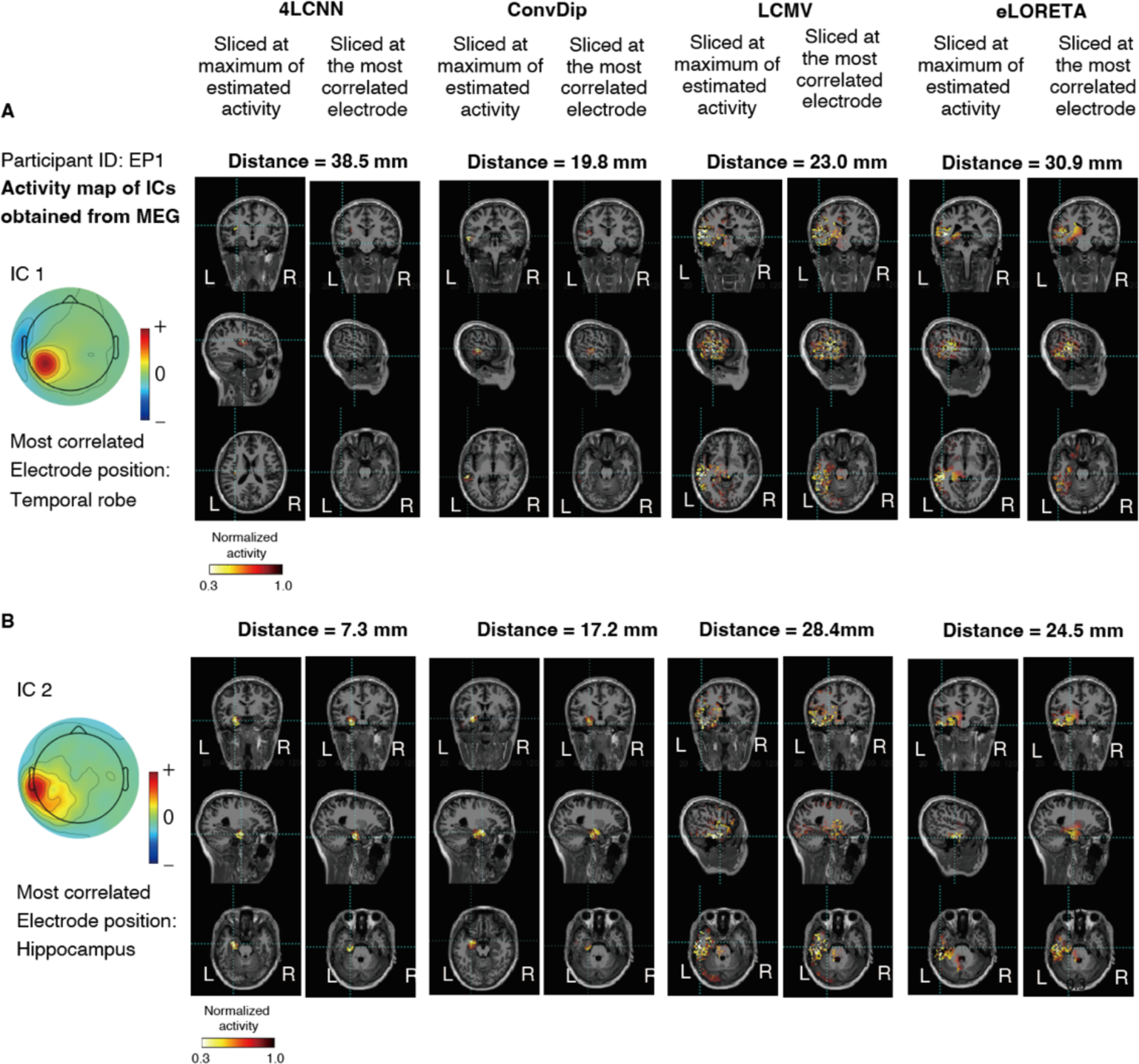
Typical examples of estimated source activity from independent components (ICs) obtained from MEG signals. For a participant (EP1), the most correlated electrodes with the ICs were located in the left temporal lobe (an ECoG electrode) and the left hippocampus (a SEEG electrode), respectively. Estimated activities of the ICs are shown in two sets of MRI views. MRI views are sliced at the most correlated electrode and an estimated source having the maximum value in the target ESI method respectively.

From the three participants, 14 ICs with correlated invasive electrodes were identified. The distances between the maximum activity locations of the estimated source activities and the corresponding most correlated electrodes did not show significant differences among the four ESI methods (4LCNN vs. ConvDip: p = 0.51, ES = 0.15; 4LCNN vs. LCMV: p = 0.51, ES = 0.28; 4LCNN vs. eLORETA: p = 0.51, ES = 0.48; ConvDip vs. LCMV: p = 0.51, ES = 0.10; ConvDip vs. eLORETA: p = 0.51, ES = 0.27; LCMV vs. eLORETA: p = 0.51, ES = 0.21) (Fig. 12A). Although the mean distances were not different among the methods, focusing on sources which were estimated spatially close to the correlated electrodes by the 4LCNN, sources with the correlated electrodes placed in the deep brain, cuneiform nucleus, and frontal pole were accurately estimated (distance ranged from 7.3 to 10.1 mm). These accurately estimated regions were relatively distant from the craniotomy site used for implanting the invasive electrodes. Spatial dispersion of the estimated source activity was also evaluated (Fig. 12B). The spatial dispersion in 4LCNN and ConvDip was significantly lower than in LCMV and eLORETA (4LCNN vs. ConvDip: p = 0.23, ES = 0.43; 4LCNN vs. LCMV: p < 0.001, ES = 1.97; 4LCNN vs. eLORETA: p < 0.001, ES = 1.61; ConvDip vs. LCMV: p < 0.001, ES = 2.01; ConvDip vs. eLORETA: p = 0.0030, ES = 1.46; LCMV vs. eLORETA: p = 0.76, ES = 0.14).

**Figure 12.**
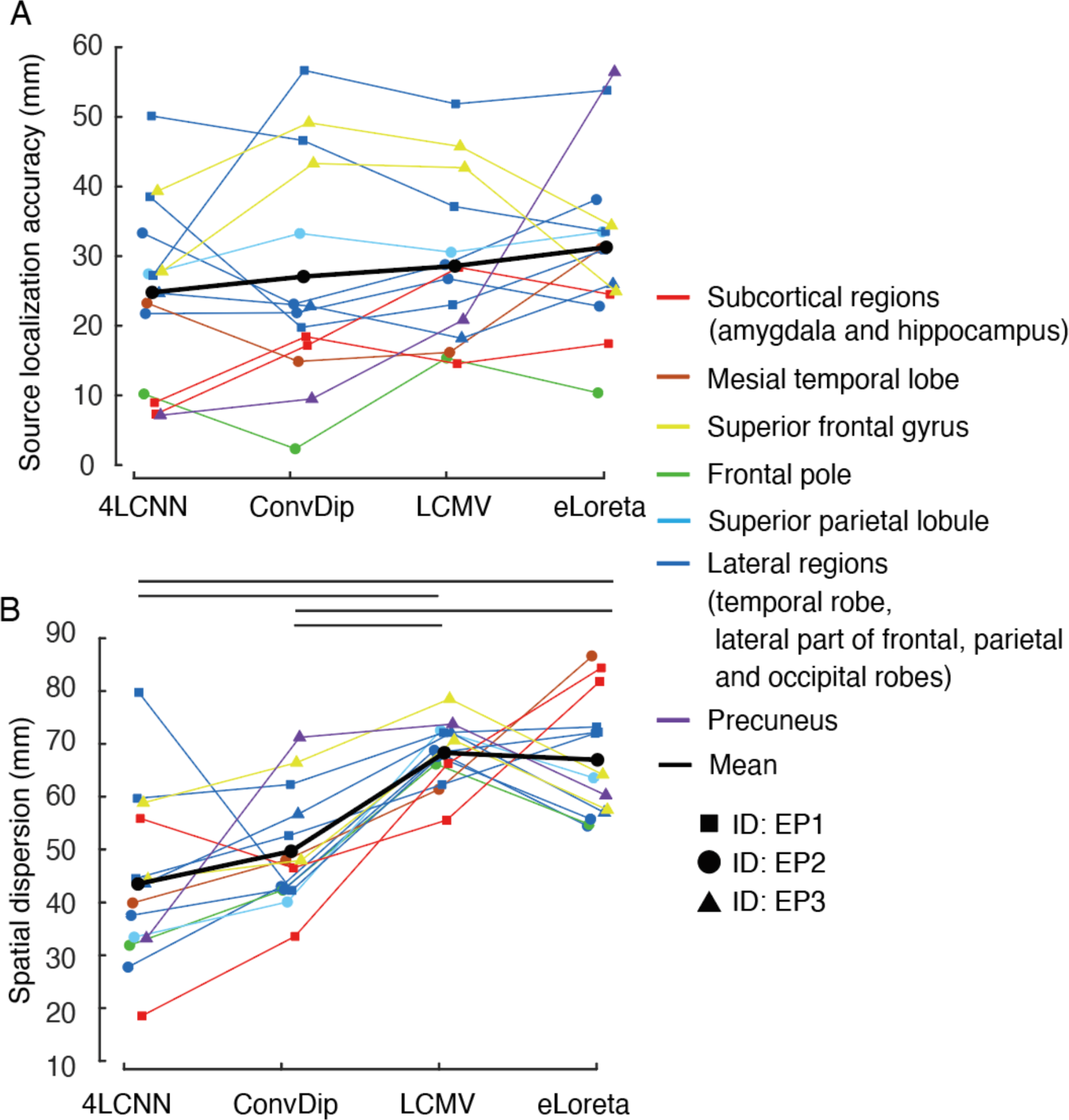
Source estimation results of independent components obtained from resting state MEG signals. (A): Source localization accuracy. (B): Spatial dispersion of the estimated source activity. (A and B): Data corresponding to each participant and brain region are presented with different markers and colors, respectively. The horizontal bars indicate a significant difference (p < 0.05, the permutation test with FDR correction).

## Discussion

The inverse problem of M/EEG source reconstruction, inherently highly ill-posed, has been tackled with our 4LCNN, which leverages realistic head models for estimating cortical and subcortical activity from M/EEG data. Compared to ConvDip, eLORETA, and LCMV, 4LCNN outperformed them in various metrics through M/EEG simulation. Its effectiveness was also confirmed by its generalizability to experimental data, aligning with somatosensory pathways and invasive electrode recordings. Although its localization accuracy was comparable to other methods in datasets with concurrent invasive recordings, this might be due to MRI’s limitations in accounting for ECoG-related brain deformations post-electrode implantation, potentially undervaluing 4LCNN’s performance (Mercier et al., 2022). Our findings highlight 4LCNN’s potential as a highly accurate method for source localization for both cortical and subcortical sources, suggesting future contributions in various neurological applications, from diagnostics to understanding cognitive and motor functions (Asadzadeh et al., 2020).

Because of the high ill-posed nature of the inverse problem of M/EEG source reconstruction, most of commonly used ESI methods require a priori assumptions to obtain a unique solution by explicitly setting regularization terms. Given the complex nature of brain activity, it is quite a challenge to set optimal regularization priors (Asadzadeh et al., 2020). Even widely utilized LORETA variants, despite their effectiveness, can lead to diffused estimations of activity (Hecker et al., 2021; Hauk et al., 2022; Sun et al., 2022). A previous study comparing different ESI methods with various regularization priors indicated that choosing between these methods involves a trade-off: smaller localization errors (e.g., sLORETA) versus reduced spatial dispersion (e.g., MNE) (Hauk et al., 2011). Therefore, setting optimal regularization priors that adequately reflect the features of M/EEG sources is a complex task.

Recent approaches have deployed artificial neural networks (ANNs) to surmount this inverse problem, adopting a data-driven approach that dispenses with regularization priors (Hecker et al., 2021; Jiao et al., 2022; Sun et al., 2022). For instance, Hecker et al. (2021) showed that a shallow CNN-based ESI method, ConvDip, could accurately estimate source localizations than conventional ESI methods. However, the success of ANN-based ESI methods has so far been demonstrated only for cortical sources (Hecker et al., 2021; Jiao et al., 2022; Sun et al., 2022). Therefore, we explored the applicability of ANN-based ESI to both cortical and subcortical sources using the 4LCNN, a model inspired by ConvDip but enhanced to augment its expressive power.

In this study, 4LCNN showed the best performance across all four ESI methods used. Regarding the localization error for maximum value sources (Figs. 6 and S9), mean errors were below 10 mm for SNRs higher than 10 dB for both cortical and subcortical sources. Remarkably, errors were as low as approximately 6.6 mm and 5.9 mm for cortical and subcortical sources, respectively, at an SNR of 30dB. The model’s robustness to noise and its capacity to focus estimations more precisely than other ESI methods underscore its potential. This accurate estimation may be attributed to the nonlinearity introduced by ReLU layers (Talathi and Vartak, 2015). Additionally, the augmented number of learnable parameters, due to enhanced input 2D topographic resolution, filters, and layers in 4LCNN compared to the preceding CNN model (ConvDip, Hecker et al., 2021), likely contributed to the improved accuracy in source localization estimation. Notably, even though the 4LCNN was trained with 20 dB SNR data, it can accurately localize EEG data across a wide range of SNR levels. Such robustness to noise has also been reported in other ANN-based ESI methods(Hecker et al., 2021; Jiao et al., 2022). The generalizability can be attributed to batch normalization (Ioffe and Szegedy, 2015) and dropout layers (Park and Kwak, 2017).

The 4LCNN estimates source activities more focally than other ESI methods (Fig. 6B). For instance, in a representative example (Fig. 4A), 4LCNN could distinguish activities between the left and right regions within the medial part of the superior frontal gyrus—a task that poses challenges for conventional ESI methods like eLORETA and LCMV. Conversely, ConvDip and 4LCNN achieved more focal estimations of source activities. This increased focality, first noted in studies introducing ConvDip. Such high focality probably stems from high model’s expressive power, which is linked to its nonlinearity and complexity. While both ConvDip and 4LCNN benefit from the nonlinearity provided by ReLU layers, the 4LCNN is a more complicated architecture. Thus, 4LCNN provides more focal estimations, even when trained on the same M/EEG simulation data as ConvDip. The estimates derived by 4LCNN were not just focal, but were able to follow source extent that was impossible with other ESI methods. (Figs. 7 and S10).

The effectiveness of neural networks in ESI tasks heavily relies on the traits of the training data, making the M/EEG simulation’s modeling a critical step for the successful application of ANN-based ESI models to actual M/EEG data. To this end, we employed the state-of-the-art brain segmentation method, CHARM, which automatically generates a highly accurate segmented MRI with delineation of ten different tissue types (Puonti et al., 2020). Additionally, we incorporated MR-compatible fiducial markers to precisely co-register MRI and EEG data. Such efforts for making precise head model prevents deviations between simulation-generated training data and actual measured M/EEG data, and is thought to contribute to the generalization performance of the created 4LCNN to real M/EEG data.

Validating 4LCNN with SEP data (Figs. 9, S7, and S8) revealed its ability to accurately localize sources in line with established neural pathways of the somatosensory system (Cruccu et al., 2008). Invasive depth electrode recordings have provided evidence that somatosensory information relays the medulla and thalamus from 14–16 ms, 16–18 ms, respectively (Macerollo et al., 2018). The 14ms component showed localized activity within the lower brainstem (medulla), although lateralization remained undetected by the 4LCNN. The 17ms component was localized in the left thalamus for two participants (HY1 and HY2), while the third participant (HY3) also exhibited significant activity in this region, albeit with the peak activity detected in the nearby left caudate. The somatosensory information by the median nerve stimulation reaches around 20 ms poststimulation (Macerollo et al., 2018).. The 4LCNN successfully estimated high activity in the somatosensory cortex’s hand area (Blankenburg et al., 2003) for all participants for the 20ms component. Thus, the 4LCNN, although trained by simulated data, is capable of providing highly accurate estimations of source locations even when estimating activities in real EEG data.

When applying 4LCNN-based ESI to time-series activities in M/EEG data that involve complex overlapping from multiple sources, it is challenging to directly input the raw signals into the 4LCNN, as the model is trained to localize a single source. Hence, we employed ICA to separate the M/EEG data into distinct signal sources (Viola et al., 2010) and assessed the source localization accuracy with ESI methods. The mean location accuracy estimated by the 4LCNN was the most precise at 24.1 mm; however, it was not significantly different from the other ESI methods (Fig. 12A). Possible reason for the lack of superiority of the 4LCNN in this validation is brain deformations resulting from the implantation of ECoG electrodes. Such implantations can lead to deformations due to electrode thickness, hematoma, tissue swelling, causing unpredictable and non-uniform brain shifts of up to 24 mm in cortical areas and up to 3 mm in deeper structures (Mercier et al., 2022). The head models for ESI were constructed from pre-implantation MRI data, because implanted electrodes significantly distort MRI data. Structural differences between the created head model and the head for which the signal was measured (an example shown in Fig. S11) potentially declined ESI accuracy, probably masking the differences in estimation accuracy among the ESI methods. Notably, among the four sources that were precisely localized by the 4LCNN (Fig. 12A), they were located deeper or far from the craniotomy site, where less deformation is anticipated. Although this finding is tentative, it suggests the precise localization capabilities of the 4LCNN when an ideal head model is used. To further validate the 4LCNN, datasets involving simultaneous SEEG and MEG recordings without ECoG would be preferable, because SEEG typically does not result in substantial brain deformation (Elias et al., 2007). In previous research employing the LAURA method, an advanced form of MNE, for ESI from concurrent SEEG and EEG recordings, the source localization error ranged between 14.8 and 23.5 mm. In comparison, the accuracy of the 4LCNN for the aforementioned four sources, which are likely less impacted by brain deformation, was superior (accuracy ranging between 7.3 to 10.1 mm). Nevertheless, additional validation is necessary to further prove the precise localization capabilities of 4LCNN due to the limited number of sources and differing experimental conditions.

In conclusion, we have proposed a data-driven ESI framework with the 4LCNN. The results presented from a series of accuracy validations involving both numerical simulations and real human data demonstrate the high feasibility of the 4LCNN for accurate M/EEG source localization of both cortical and subcortical sources. The 4LCNN’s highly accurate source localization throughout the entire brain will significantly contribute to the clinical diagnosis and understanding of the pathophysiology of various diseases and brain functions.

## Supplementary figures

**Figure S1:**
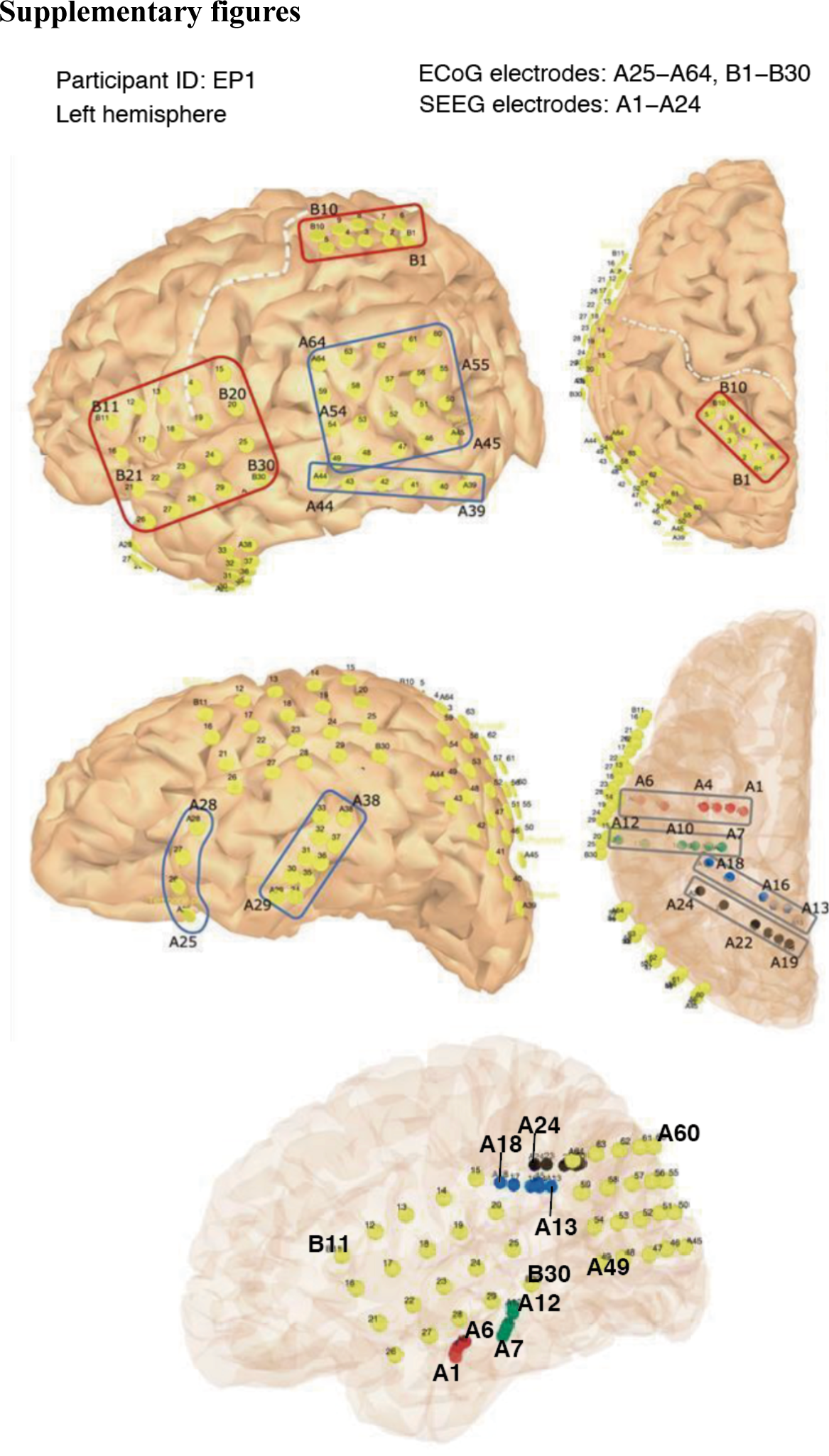
Placement of invasive electrodes in a participant (EP1). Locations of ECoG electrodes are shown with yellow circles. Locations of SEEG electrodes are indicated in colors other than yellow.

**Figure S2:**
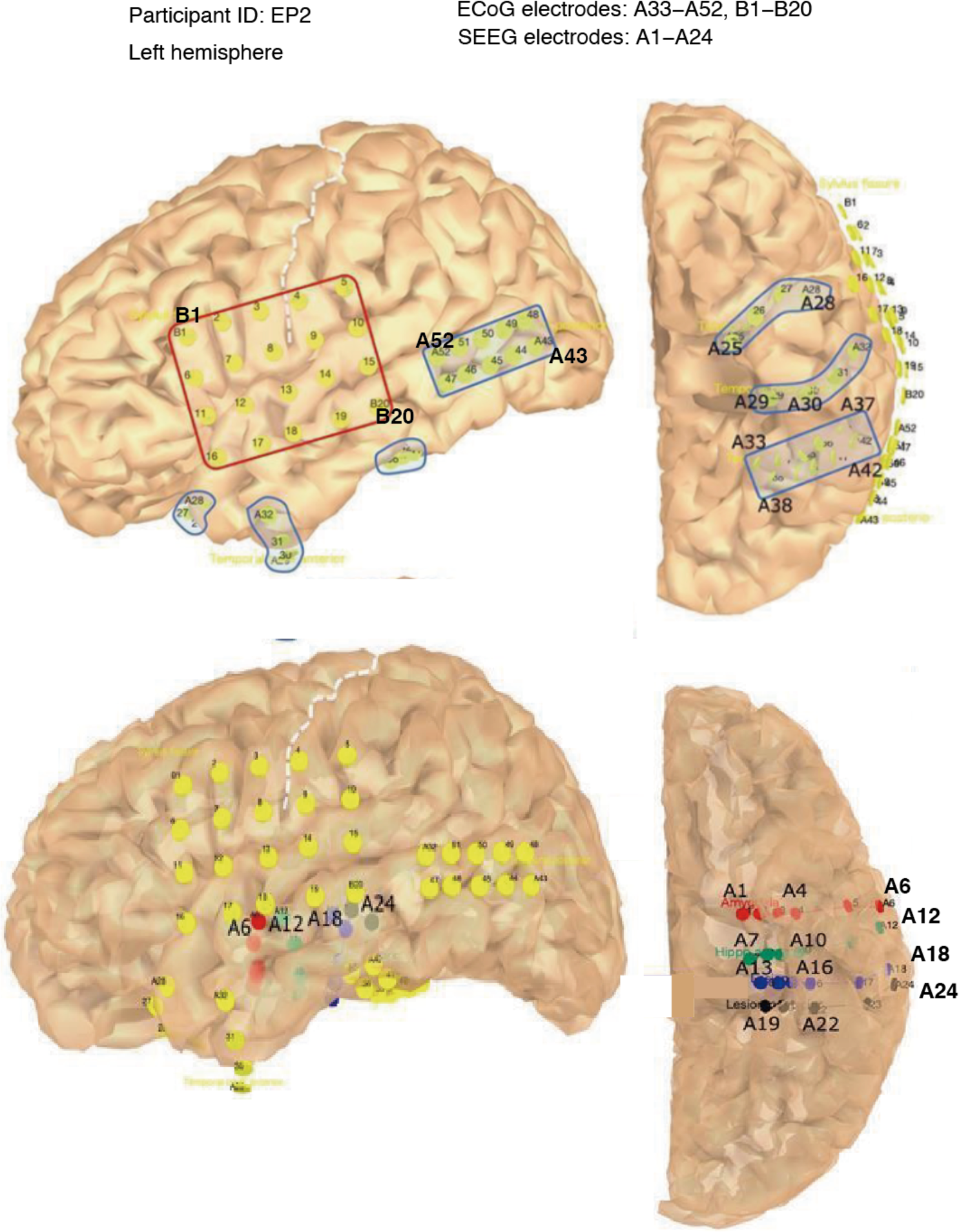
Placement of invasive electrodes in a participant (EP2). Locations of ECoG electrodes are shown with yellow circles. Locations of SEEG electrodes are indicated in colors other than yellow.

**Figure S3:**
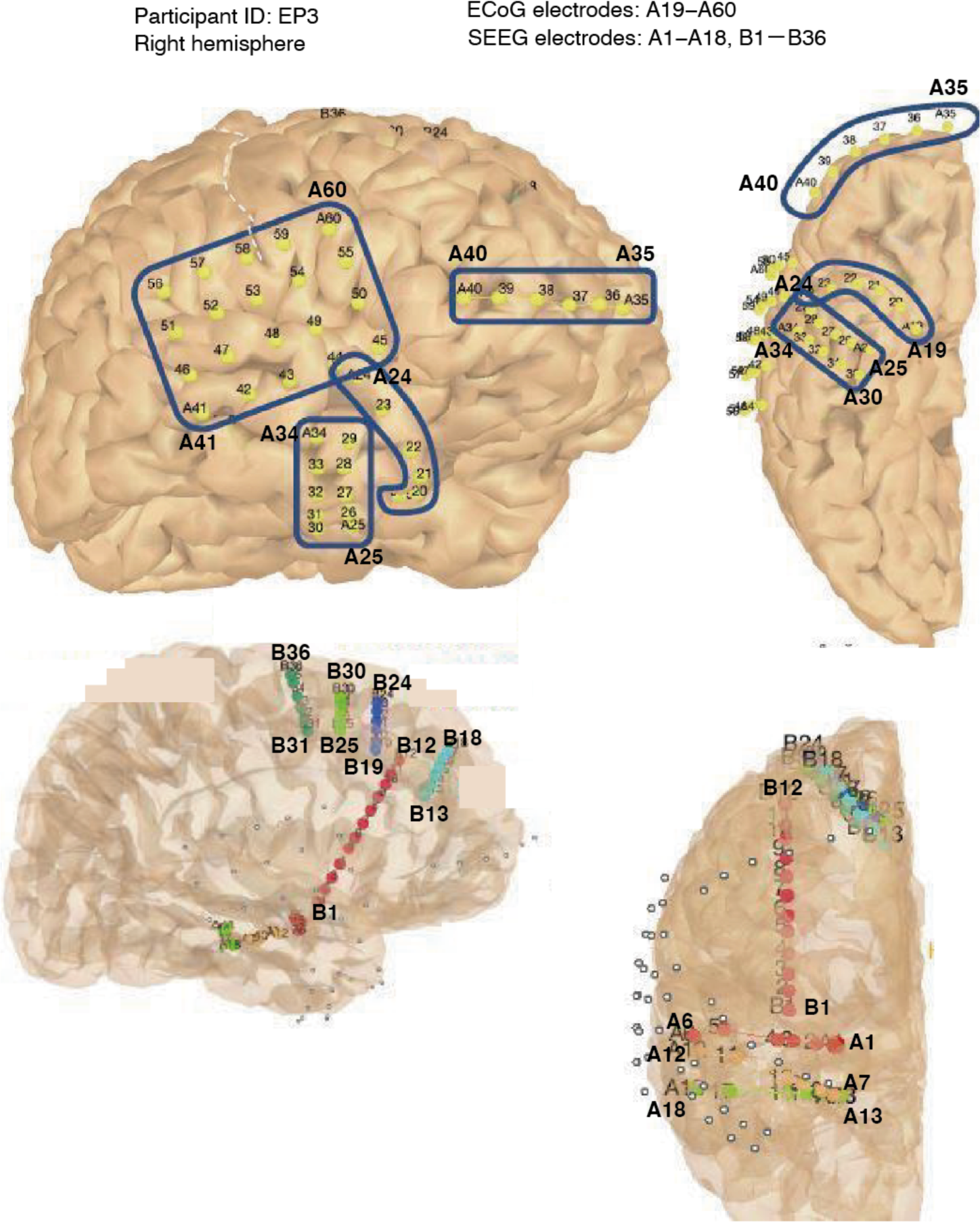
Placement of invasive electrodes in a participant (EP3). Locations of ECoG electrodes are shown with yellow circles. Locations of SEEG electrodes are indicated in colors other than yellow.

**Figure S4.**
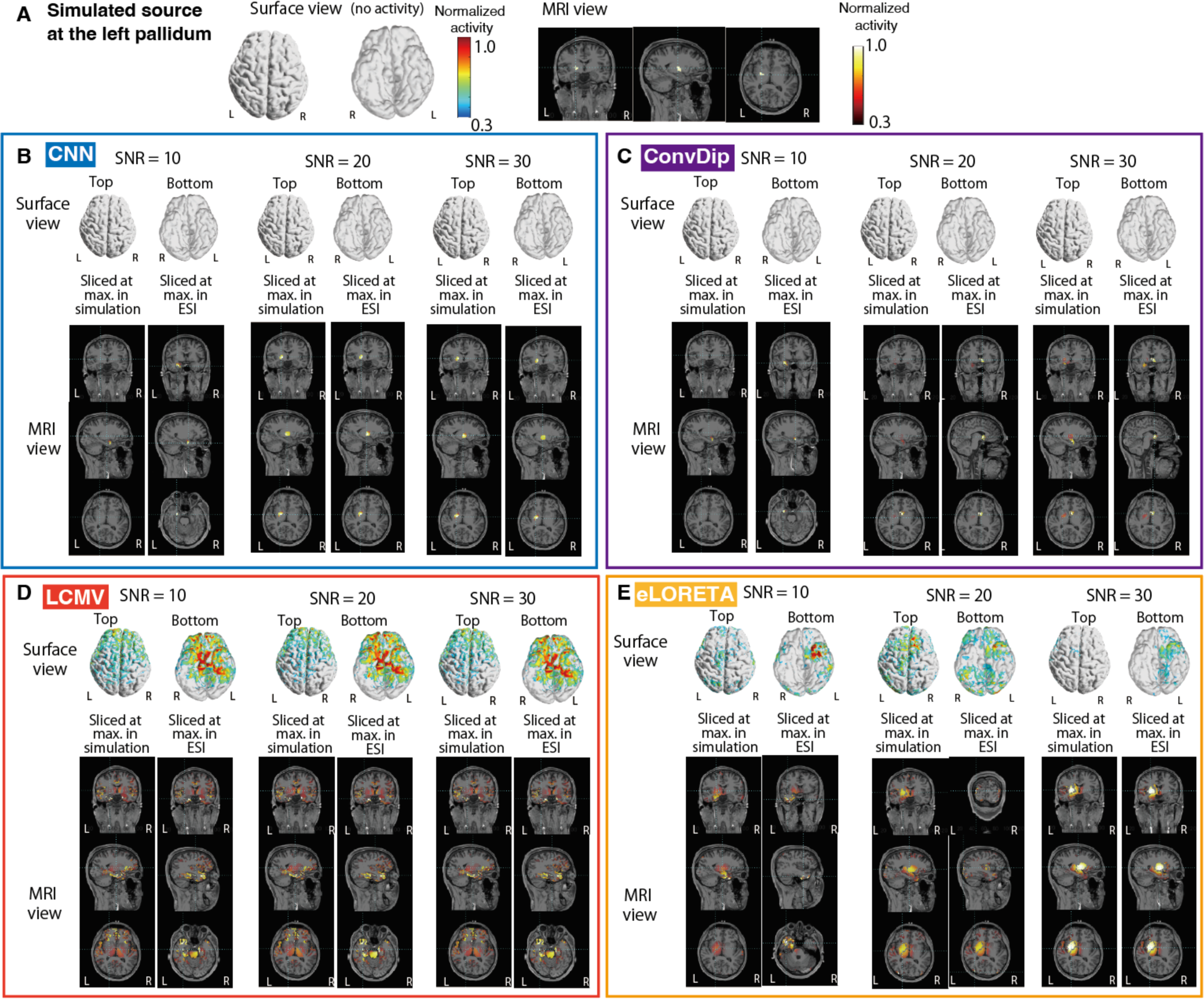
Examples of estimated subcortical source activity located at the brainstem (the lower pons). (A): Simulated source activity is shown in surface and MRI views. There is no activity in the surface view since this example is hippocampus source. (B‒E): Estimated source activity by 4LCNN, COnvDip, LCMV, and eLoreta are shown in (B), (C), (D) and (E), respectively. Different levels of noise were added in EEG simulations. Results of estimation for three different SNR conditions (SNR = 10, 20, and 30) are shown. The activities are shown in as surface and MRI views. MRI views are sliced at two different locations: sources with the maximum values in the simulated sources and the target method, respectively.

**Figure S5.**
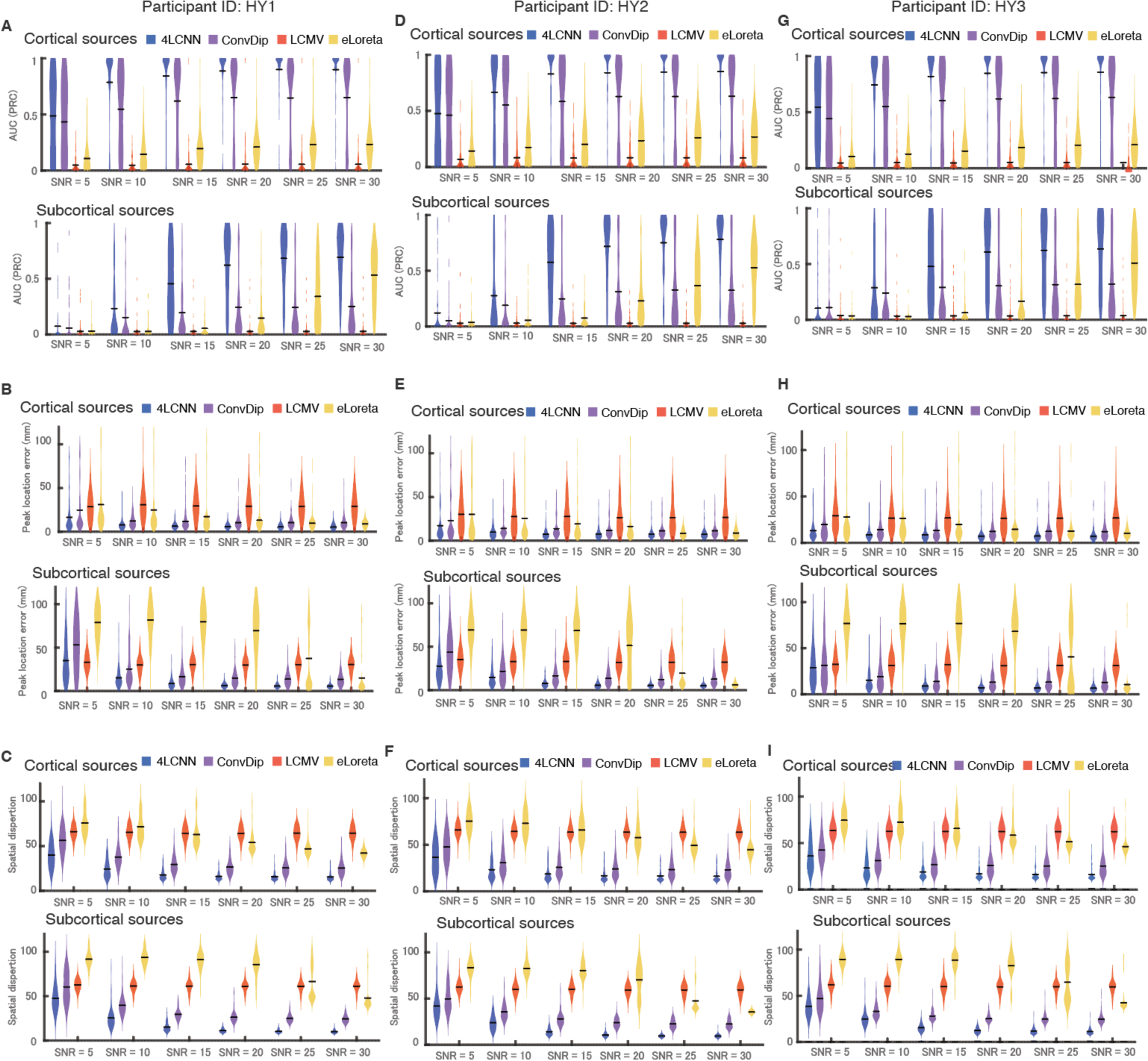
Individual data (HY1, HY2 and HY3) of three metrics for evaluation of estimation accuracy in simulation data. Violin plots of the distributions of three different metrics (area under the Precision-Recall curve (AUPRC), localization error of peak activity, and spatial dispersion) of 100 simulation data are shown. The bars in the violin plots indicate the mean values.

**Figure S6.**
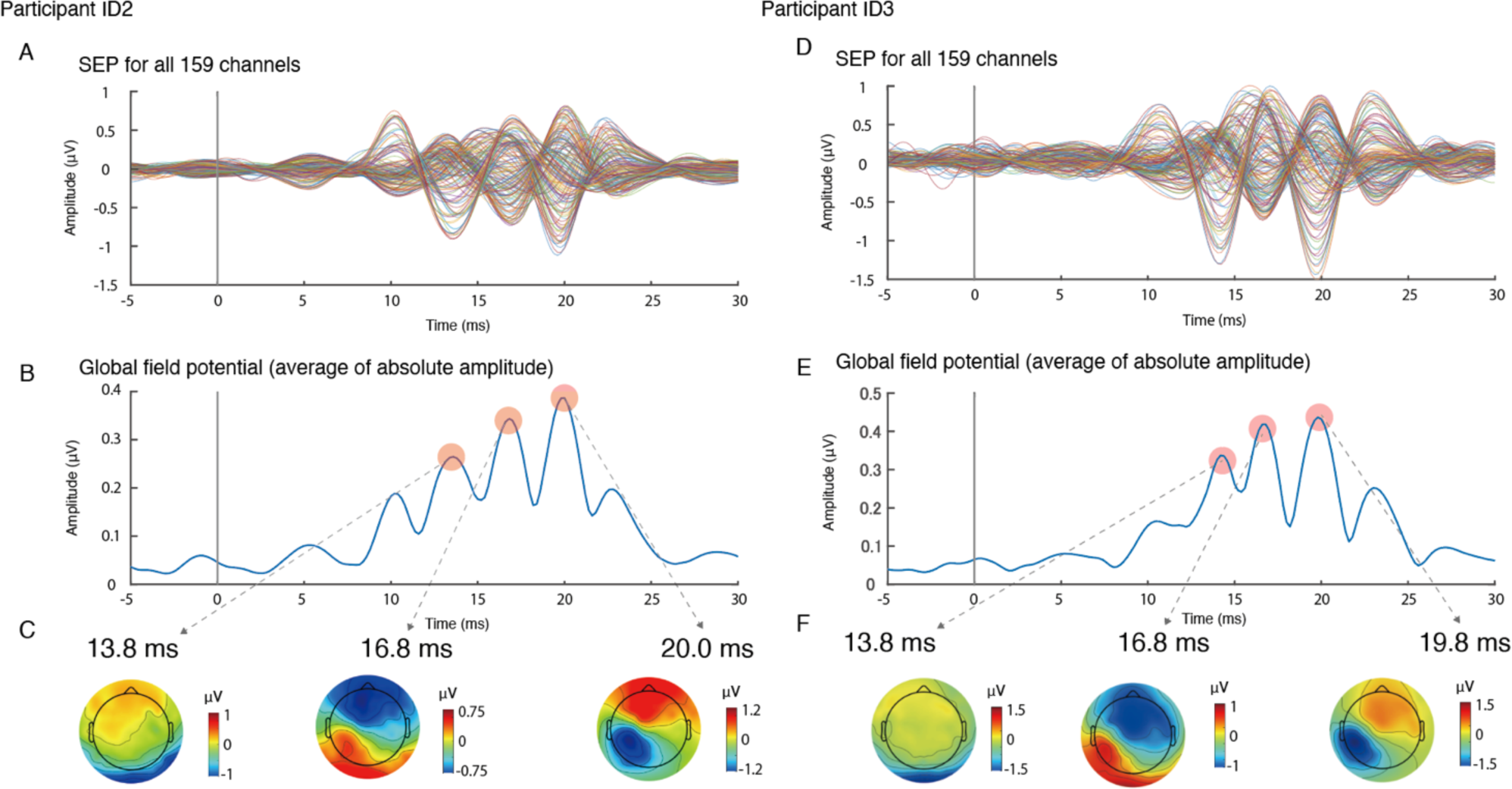
Somatosensory evoked potentials (SEPs) in HY2 and HY3. (A and D): An example of obtained SEPs in all 159 channels. (B and E): Global field potentials, which is the average of absolute amplitude across all the channels. There are three clear peaks corresponding to the components illustrated in (A/D). (C and F): 2D topographic maps for the peak activities.

**Figure S7.**
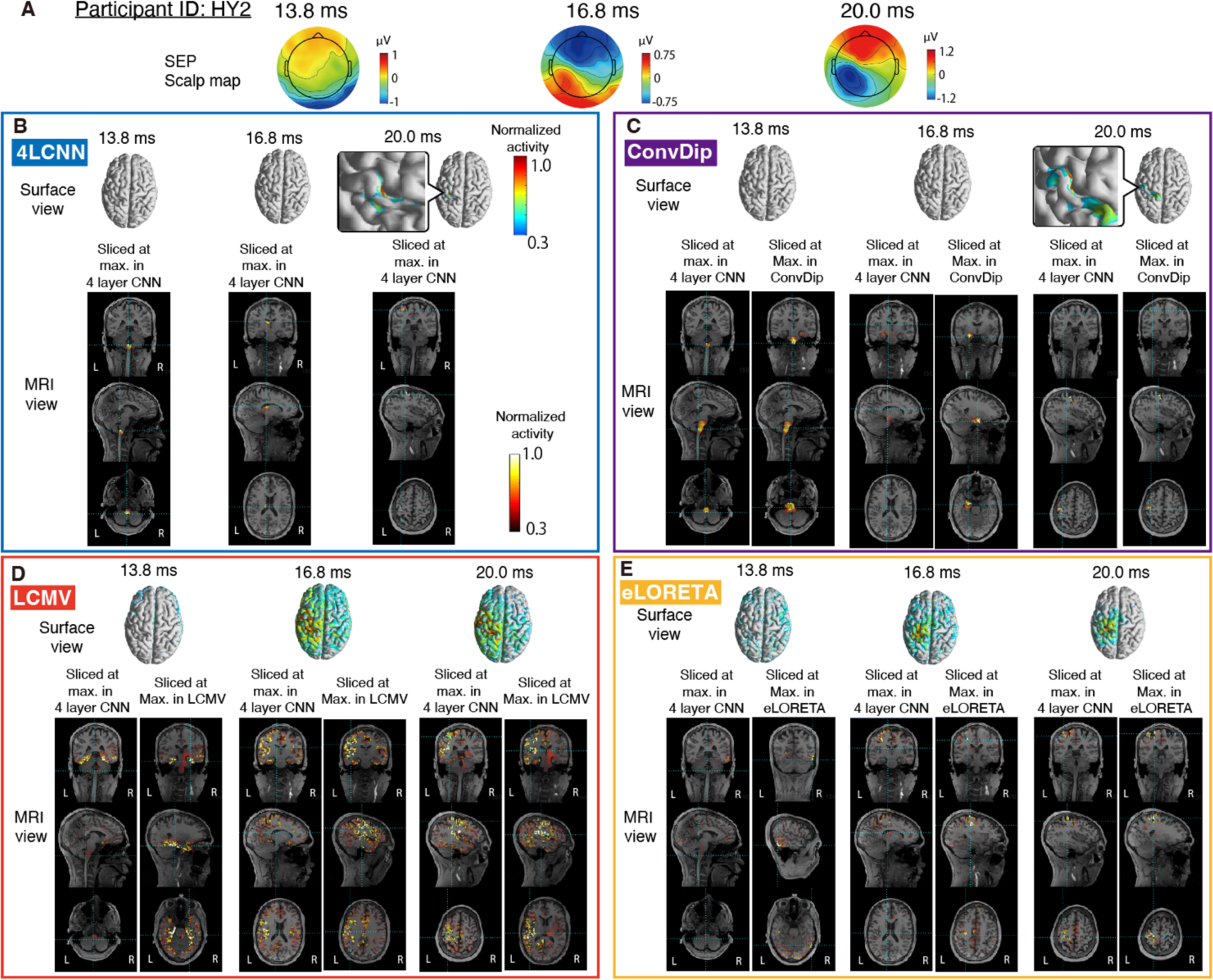
Source activity estimation of SEP in a participant (participant ID: HY2). (A): 2D topographic maps for three peak activities of SEP. (B‒D): Estimated source activity are shown in surface and MRI views. (B): Estimated source activity by 4LCNN is displayed. MRI views are sliced at the source corresponding to the maximum value. (C−E): Estimated source activity by ConvDip(B), LCMV (C), and eLORETA (E). Two sets of MRI views are shown. MRI views are sliced at sources with the maximum values in 4LCNN and the target method respectively.

**Figure S8.**
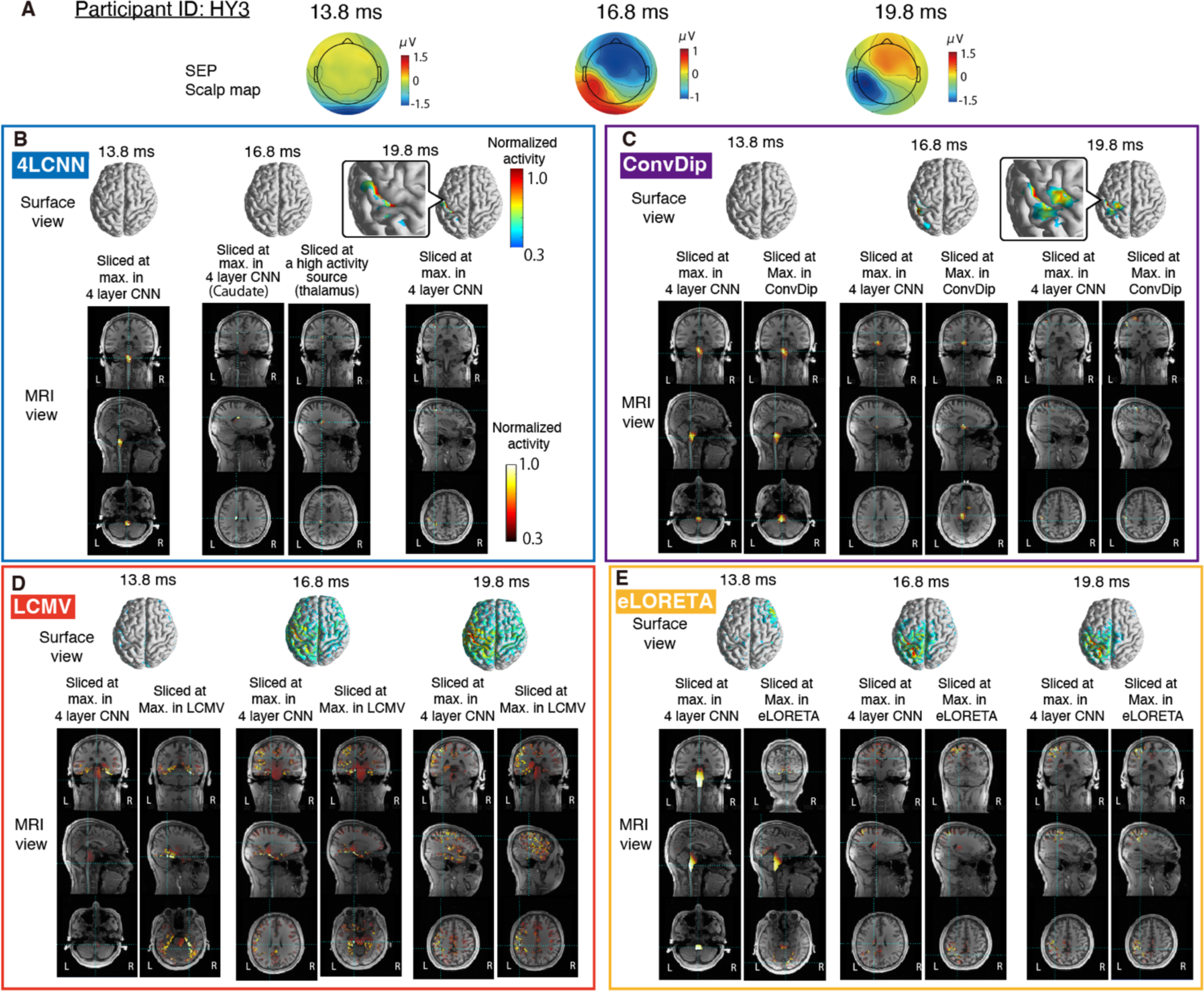
Source activity estimation of SEP in a participant (participant ID: HY3). (A): 2D topographic maps for three peak activities of SEP. (B‒D): Estimated source activity are shown in surface and MRI views. (B): Estimated source activity by 4LCNN is displayed. MRI views are sliced at the source corresponding to the maximum value across all the sources. The maximum value was estimated in the caudate. The MRI view is also sliced at the thalamus, which also showed high value. (C−E): Estimated source activity by ConvDip (B), LCMV (C) and eLoreta (D). Two sets of MRI views are shown. MRI views are sliced at sources with the maximum values in 4LCNN and the target method respectively.

**Figure S9.**
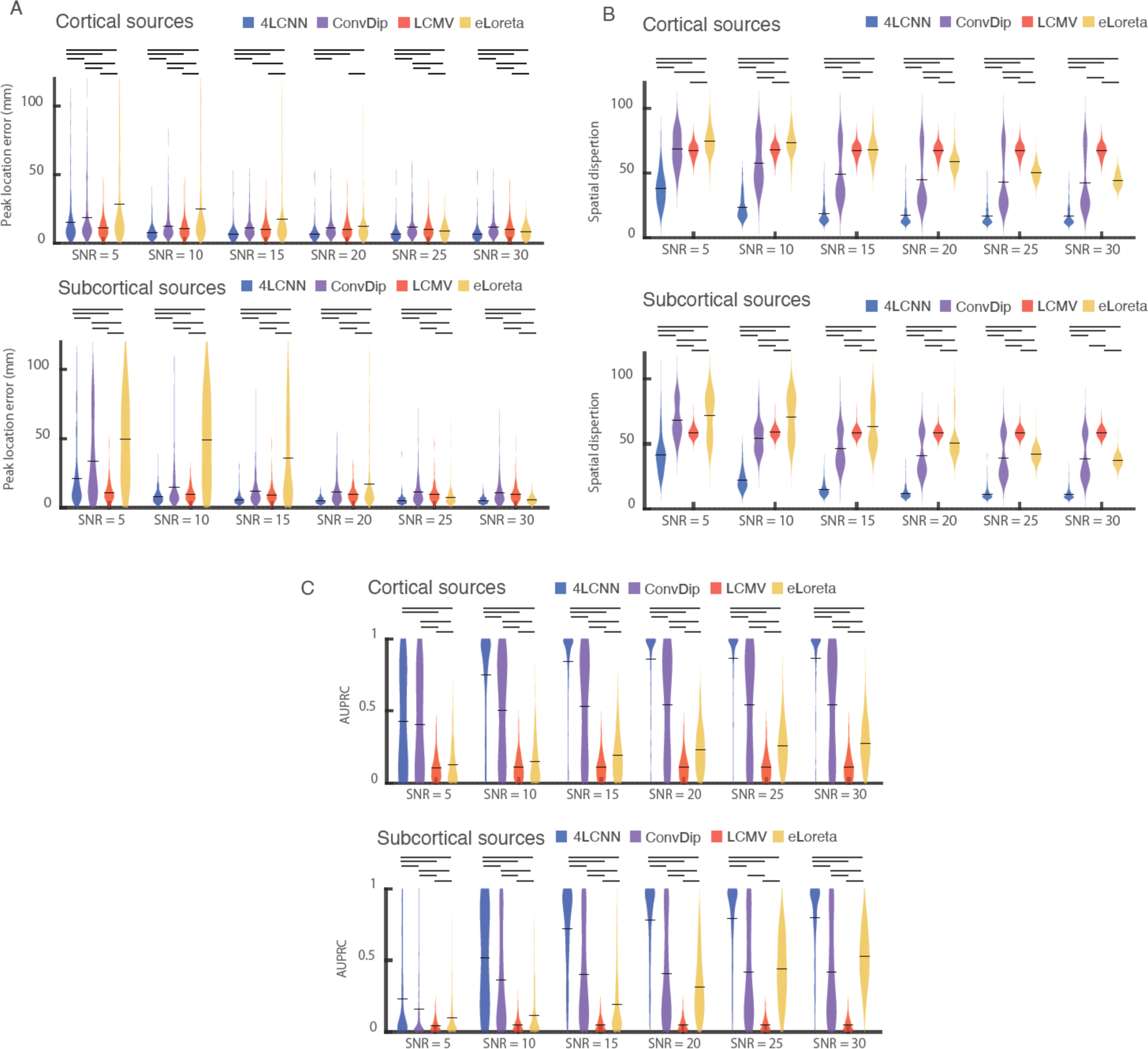
Quantitative evaluation of estimation accuracy in simulation data for MEG. Violin plots of the distributions of three different metrics (area under the Precision-Recall curve (AUPRC), localization error of peak activity, and spatial dispersion) of 300 simulation data from three participant head models (a hundred for each) are shown in (A), (B) and (C), respectively. The bars in the violin plots indicate the mean values. The bars above the violin plots indicate significant differences between conditions (p < 0.05 by the permutation test with FDR correction).

**Figure S10.**
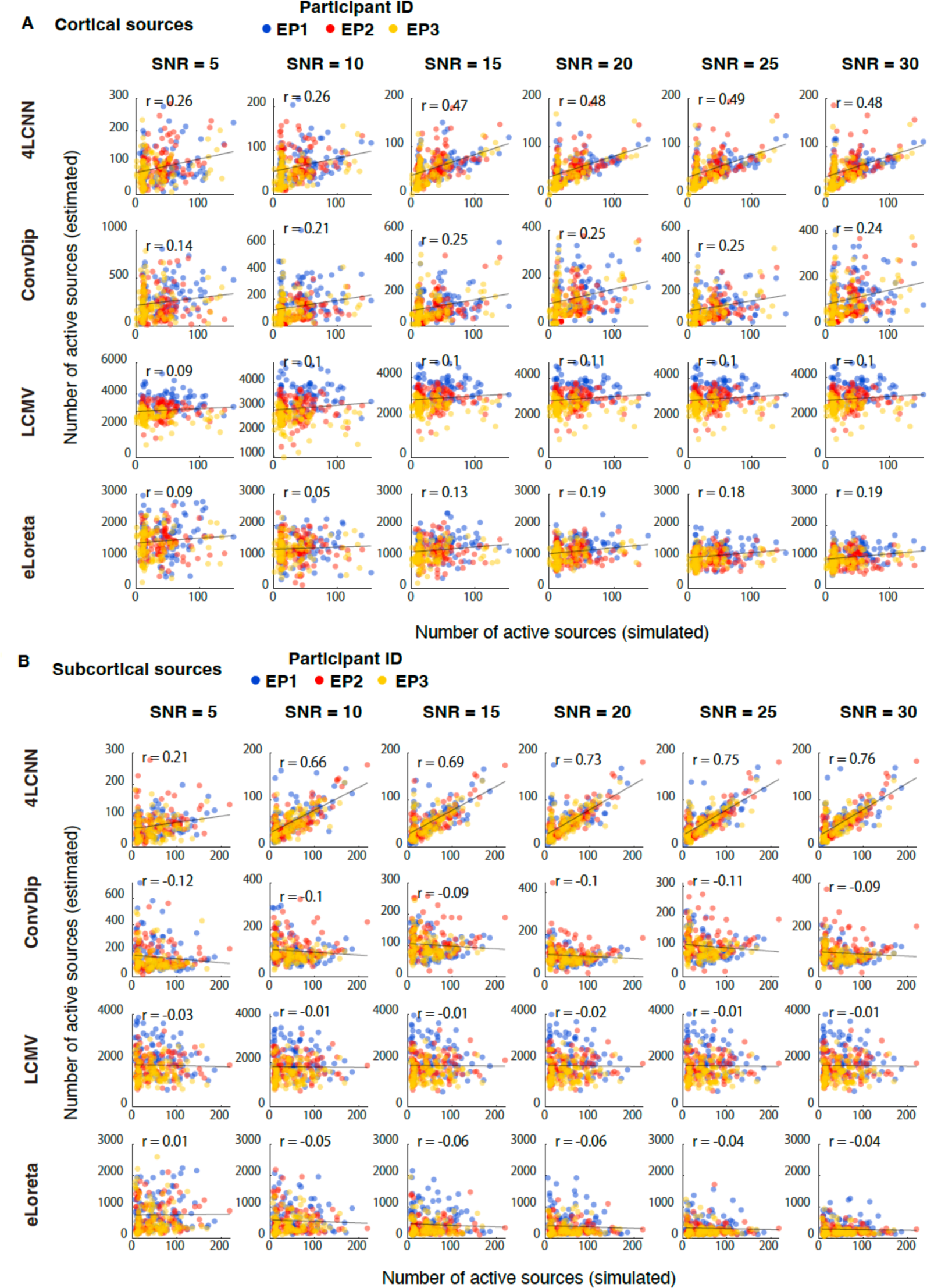
Correlation of activation area (i.e., number of active sources) between simulated and estimated source activities of 300 simulation of MEG data from three participant head models for cortical source simulation (A) and subcortical source simulation data(B). Each participant data is plotted as closed circles in a different color.

**Figure S11.**
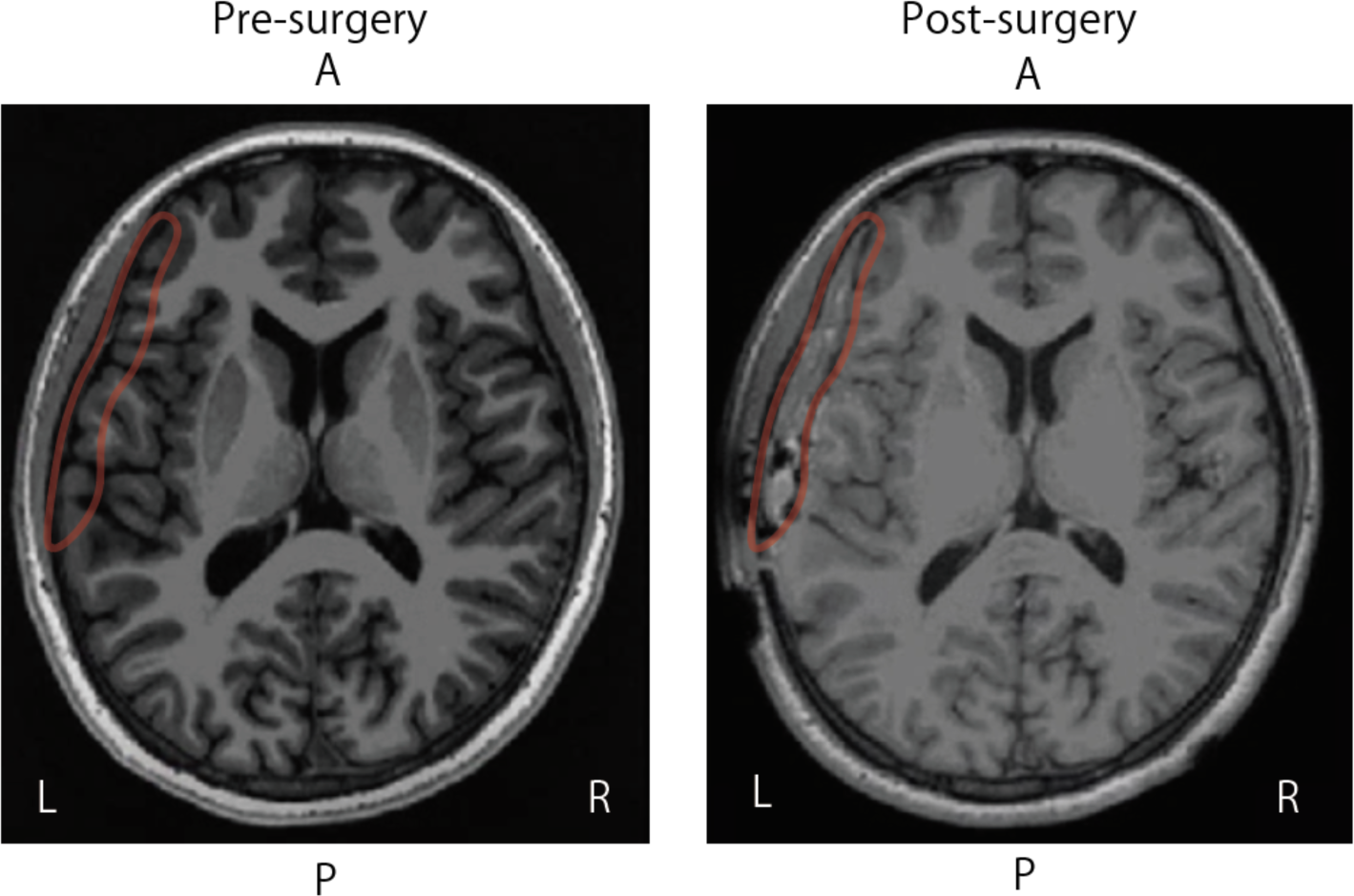
An example of deformation of the brain after implantation of invasive electrodes from a participant (EP1). (A) A T1 weighted MRI pre-electrode implantation. (B) A T1 weighted MRI post-electrode implantation. The area supposed to be hematoma in the post-surgery image is highlighted by a red line. The same region is also highlighted in the pre-surgery image.

